# BenchXAI: Comprehensive Benchmarking of Post-hoc Explainable AI Methods on Multi-Modal Biomedical Data

**DOI:** 10.1101/2024.12.20.629677

**Authors:** Jacqueline Michelle Metsch, Anne-Christin Hauschild

## Abstract

The increasing digitalisation of multi-modal data in medicine and novel artificial intelligence (AI) algorithms opens up a large number of opportunities for predictive models. In particular, deep learning models show great performance in the medical field. A major limitation of such powerful but complex models originates from their ’black-box’ nature. Recently, a variety of explainable AI (XAI) methods have been introduced to address this lack of transparency and trust in medical AI. However, the majority of such methods have solely been evaluated on single data modalities. Meanwhile, with the increasing number of XAI methods, integrative XAI frameworks and benchmarks are essential to compare their performance on different tasks. For that reason, we developed BenchXAI, a novel XAI benchmarking package supporting comprehensive evaluation of fifteen XAI methods, investigating their robustness, suitability, and limitations in biomedical data. We employed BenchXAI to validate these methods in three common biomedical tasks, namely clinical data, medical image and signal data, and biomolecular data. Our newly designed sample-wise normalisation approach for post-hoc XAI methods enables the statistical evaluation and visualisation of performance and robustness. We found that the XAI methods Integrated Gradients, DeepLift, DeepLiftShap, and GradientShap performed well over all three tasks, while methods like Deconvolution, Guided Backpropagation, and LRP-*α*1-*β*0 struggled for some tasks. With acts such as the EU AI Act the application of XAI in the biomedical domain becomes more and more essential. Our evaluation study represents a first step toward verifying the suitability of different XAI methods for various medical domains.

## 1 Introduction

In recent years, Machine Learning (ML) and particularly Deep Neural Networks (DNNs) have shown great potential in biomedical applications such as medical imaging, genomics, and clinical data analysis. DNNs are especially effective at modeling complex, nonlinear relationships, which is often necessary when dealing with multidimensional data. Their capability to handle large and complex data has often allowed them to surpass traditional methods in terms of predictive accuracy and diagnostic performance. For instance, DNNs have achieved state-of-the-art performance in tasks like species-agnostic transfer learning [35], classification of abnormalities in 12-lead ECGs [5], and even in fields such as graph classification [31].

However, transferring these networks into clinical practice, for example in the form of clinical decision support systems (CDSS), is often difficult due to their ‘black-box’ nature. Unlike traditional statistical models, such as linear regression models, which are often interpretable by design, DNNs are highly complex networks, making it difficult to understand the rationale behind their predictions. In a setting where these networks might directly impact an individual’s health, a lack of interpretability is unacceptable [29]. Clinicians need to trust that the network’s recommendations align with clinical reasoning, which is not possible without a clear understanding of how a DNN arrives at its conclusions. These interpretability issues enforced legislative and regulatory acts that are currently being shaped in frameworks such as the EU AI Act [27].

To aid in increasing the transparency of AI, the field of explainable AI (XAI) has emerged, motivating the development of many different methods that aim to fill this gap and add global interpretability and prediction specific understanding. XAI methods are typically divided into two categories: ante-hoc and post-hoc methods. Ante-hoc methods provide explanations during the training process, e.g., global model coefficients of linear regression models or calculated gini impurity scores of random forest classifiers. In contrast, post-hoc methods are applied after training an AI model. These methods specifically focus on adding interpretability to the aforementioned ‘black-box’ models, such as DNNs, by generating primarily local explanations for the predictions of each sample. These explanations are often created by making use of the gradient which is calculated using the backpropagation algorithm used to train the DNN [51, 55] or by occluding certain parts of the input sample and analyzing the effect on the prediction [62]. Other methods, such as Shapley Values [54], make use of algorithms already known from game theory.

A big issue with these methods is the huge variety of underlying theories, objectives, and corresponding implementations. Captum AI [26], one of the most widely used and comprehensive platforms, supports 16 different XAI methods. Additionally, a major difficulty is that there is no real consensus on which methods to use in which use case or data modality. This is mostly because it is hard to validate the explanations generated by these methods. Therefore, the first evaluations were mostly done on image data sets where explanations could be directly validated on known pixel based ‘gold-standards’. The explanations are represented in the form of saliency maps [63] directly on top of the image showing which regions were relevant for a specific classification task. However, for other tasks, such as medical images where the importance of pixel areas is not certain, and other data modalities, such as electronic health records or gene expression data, the evaluation is more difficult. This is especially true since there is usually no ‘gold-standard’ against which to validate the explanations of XAI methods.

Due to the described challenges with respect to the evaluation and the huge variance in explanations across XAI methods, large benchmarking studies are critical to show which approach can be applied for which task. Particularly, the inconsistent explanations between XAI methods could potentially reduce trust in AI systems and thus make integration into clinical practice even harder. Moreover, in the worst case, differing explanations could also lead to the selection of the explanation that confirms the pre-existing beliefs (confirmation bias) of the user, potentially enhancing biased decision-making.

### 1.1 Literature

Currently, only a few benchmark studies have been published to address this, yet these either focus solely on implementations of evaluation frameworks for XAI or on benchmarking one specific data modality. Agarwal et al. [2] introduced the OpenXAI platform, which provides an automated end-to-end pipeline that currently only supports tabular data. They also provide a short benchmark of six XAI methods on some evaluation metrics, such as stability and fairness. The Xplique platform [16] is similar to the Captum AI [26] platform but with added evaluation metrics and saliency maps. This platform is mostly implemented for image data, and no actual benchmarking results have been published. Duell et al. [13] compared three different XAI methods on one electronic health record (EHR) dataset, while Hu et al. [20] used SHAP [44] to develop a predictive model on intensive care data. On clinical imaging, Shinde et al. used CAM for pathology localization on MRI data [47], and Shad et al. used LIME [42] on MRI data for Alzheimer’s disease prediction. Bender et al. [5] compared Integrated Gradients [55] and LRP [4] on 12-lead ECG data. LIME [42] was used by Kamal et al. [23] to interpret which genes were responsible for Alzheimer’s disease in patients while Dwivedi et al. [15] used Integrated Gradients [55], GradientShap [44] and DeepLift [48] for biomarker discovery for lung cancer classification. Lastly, Srinivasu et al. [53] used SHAP [44] on their CatBoost+MLP model for breast cancer prediction to achieve transparency of their model.

### 1.2 Objectives, Contribution, and Structure of the Paper

To the best of our knowledge, there is no large evaluation study comparing the performance of a multitude of XAI methods on different data modalities. Due to the diversity of biomedical data, a large evaluation study of XAI methods across multiple biomedical data modalities is needed. We aim to address the unsolved issues of evaluating and comparing XAI methods across multiple modalities in biomedicine. The structure of the whole paper is divided into three typical biomedical modalities, namely clinical data, medical image and signal data, and molecular data (see Figure 1). The major contributions from this study are summarized below:

- Implementation of a comprehensive XAI evaluation framework called BenchXAI, comprising fifteen state-of-the-art XAI methods. Its major advantage is its flexibility in combining various machine learning methods and data modalities.
- Establishing a generalized evaluation scheme, including standardized normalisation of scores as well as statistical analysis evaluation and visualisations of performance and robustness, allowing for a comparison across ML, XAI, and data modalities.
- Evaluation of task-specific advantages and disadvantages of XAI in three typical biomedical modalities, namely clinical data, medical image and signal data, and molecular data.

**Figure 1:**
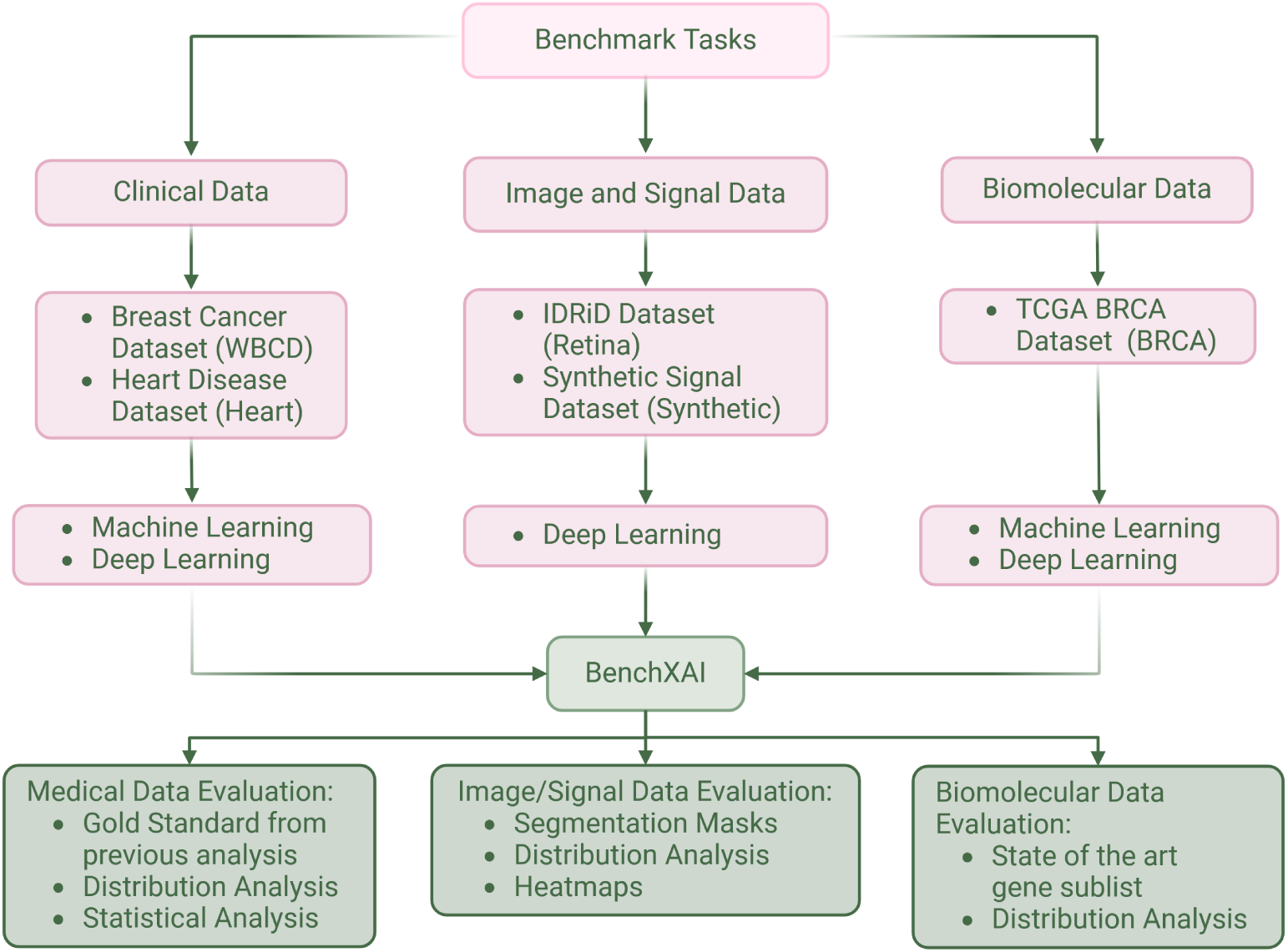
Workflow depiction showing the different benchmarking tasks. These tasks include different datasets, machine learning (ML) and Deep Learning (DL) models, our BenchXAI package for applying the different XAI methods, and different evaluations of these XAI methods.

## 2 Benchmarking Tasks and Datasets

In order to comprehensively benchmark XAI methods on all relevant biomedical data modalities, we split our evaluation into three subtasks (see Figure 1) based on the different data modalities (see Table 1) used in this study, since they had to be processed slightly differently and all come with their own specific challenges and evaluation strategies. The sections throughout the paper will all address these different subtasks.

**Table 1:** Overview over all datasets used in this study. Information on the benchmarking task, type of data, number of samples, as well as, balancedness of the binary target classes.

| Dataset Name | Task | Type | # Samples | #Features | % Class 0 | % Class 1 |
| --- | --- | --- | --- | --- | --- | --- |
| WBCD | 1 | Tabular | 699 | 9 | 65.52% | 34.48% |
| Heart | 1 | Tabular | 299 | 12 | 67.89% | 32.11% |
| Signal | 2 | Signal | 15000 | (6,500) | 52.32% | 47.68% |
| IDRiD | 2 | Image | 514 | (570,858) | 32.68% | 67.32% |
| BRCA | 3 | Tabular | 867 | 867 | 19.72% | 80.28% |

### 2.1 Task 1: XAI Benchmark for Clinical Data

Our first task will be to benchmark different XAI methods on clinical data, such as electronic health records (EHR). Clinical data often has a problem with having too few samples to train machine learning (ML) or deep learning (DL) models. Additionally, it is much harder to validate the prediction results of a DL model since there is no easily understandable visual explanation, and we often do not have a gold standard for explanations. The **Wisconsin Breast Cancer Dataset (WBCD)** [30, 58] is a dataset that comprises 699 samples taken from fine needle aspirates (FNA) from the patient’s breast. Since the presence of a breast mass does not always indicate a malignant cancer, FNA is a non-invasive and cost-effective diagnostic that can help evaluate malignancy. The data comes from the University of Wisconsin Hospital and was collected to accurately diagnose breast masses based solely on an FNA test from 1992. For this task, nine visually assessed features of an FNA sample were identified and assigned an integer value between 1 and 10, with one being closest to benign. A full list of the features can be found in the supplementary.

Many rule sets have already been reported for the WBCD dataset to increase classification accuracy [34, 12, 6, 9]. Hayashi and Nakano [18] compared these rule extraction methods using 10-fold cross validation. They also implemented their own rule extraction method, resulting in a rule set of four rules, and achieved an accuracy of 95.80%. Their rule set only uses the Bare nuclei (BN) and Clump thickness (CT) features, which is less than all the other rule sets. Our goal will be to assess whether XAI methods will find BN and CT to be the most important features or if the AI model relies on more than these two features for their predictions.

The **Heart Disease Dataset (Heart)** contains medical records of 299 patients who experienced heart failure. The dataset comprises 13 features collected at the Faisalabad Institute of Cardiology and Allied Hospital in Faisalabad-Pakistan from April to December 2015 [3]. A full list of the features can be found in the supplementary, and a detailed description of the data collection and features can be found in [3, 11]. The target for the binary classification task is the death of a patient during their follow-up period [3].

All nonbinary features were normalised using *L*_2_ normalisation. In their paper, Chicco and Jurman [11] show that the features serum creatinine (SC) and ejection fraction (EF) suffice to predict the survival of patients. In contrast, Ahmad et al. [3] in addition, identified Age, high blood pressure (HBP), and anaemia (A) as important in the survival prediction. In our study, we will use different XAI methods to validate the importance of these features.

### 2.2 Task 2: XAI Benchmark for Medical Image and Signal Data

Our second task is the benchmarking of different XAI methods on medical image and signal data, such as MRI or CT images and EEG or ECG signals. Image classification has already been readily studied using XAI methods, but medical images are often times much larger, and important details much smaller. Nonetheless, DL models oftentimes perform very well in classifying these highly complex medical images, so understanding how they reach a specific prediction is crucial for gaining more insights into how they work. For signal data, there is an added difficulty of not only spatial information but also time-dependent, sometimes periodical features, which need to be identified by a DL model.

To address the lack of a ground truth dataset for time-series classification, Turbé et al. [56] developed a **Synthetic Signal Dataset (Synthetic)** replicating the complexity of time-series classification tasks. The dataset forces the network to learn temporal dependencies and dependencies across features. The dataset comprises six features with 500 time steps. Each feature comprises a random baseline simulated by a sine wave with its amplitude multiplied by 0.5 and frequency sampled from a discrete uniform distribution ∼ U(2, 5). Two random features *f*_1_ and *f*_2_ are selected for which sine waves with a support of 100 time steps are added at random points in time. The frequencies of these sine waves are randomly sampled from a discrete uniform distribution ∼ U(10, 50). For the rest of the four features, a square wave with frequency ∼ U(10, 50) is added with a probability of 50%. The classification task will be to predict whether the sum of the two frequencies of *f*_1_ and *f*_2_ is larger than 60, which results in a balanced dataset. An exemplary sample is shown in Figure 2. Since we know exactly where the network should look for this dataset, we can calculate exactly how many XAI attributions lie inside the relevant time frames and how many lie outside.

**Figure 2:**
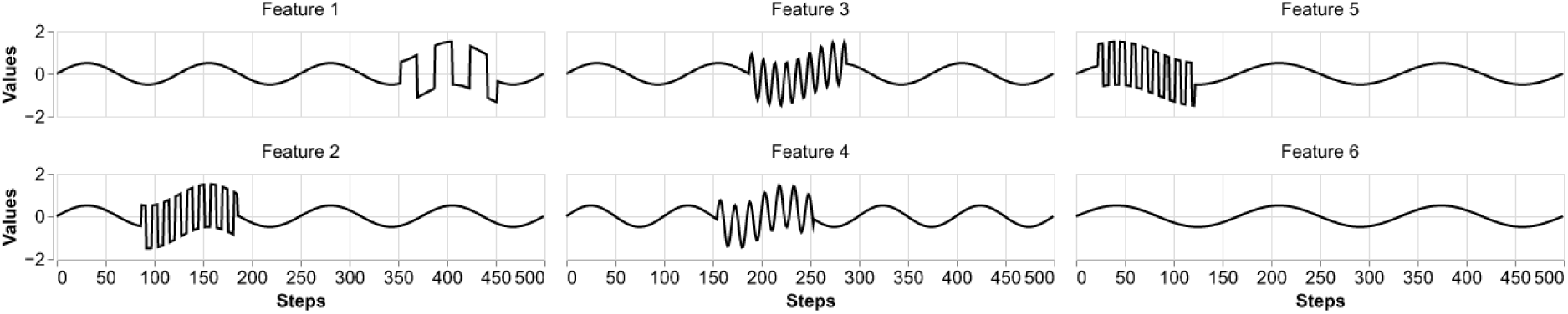
An exemplary sample from the synthetic dataset. The classification task is to predict whether the frequencies of the sine waves in features 3 and 4 are larger than 60. The square waves in features 1, 2, and 5 are noise.

**Figure 3:**
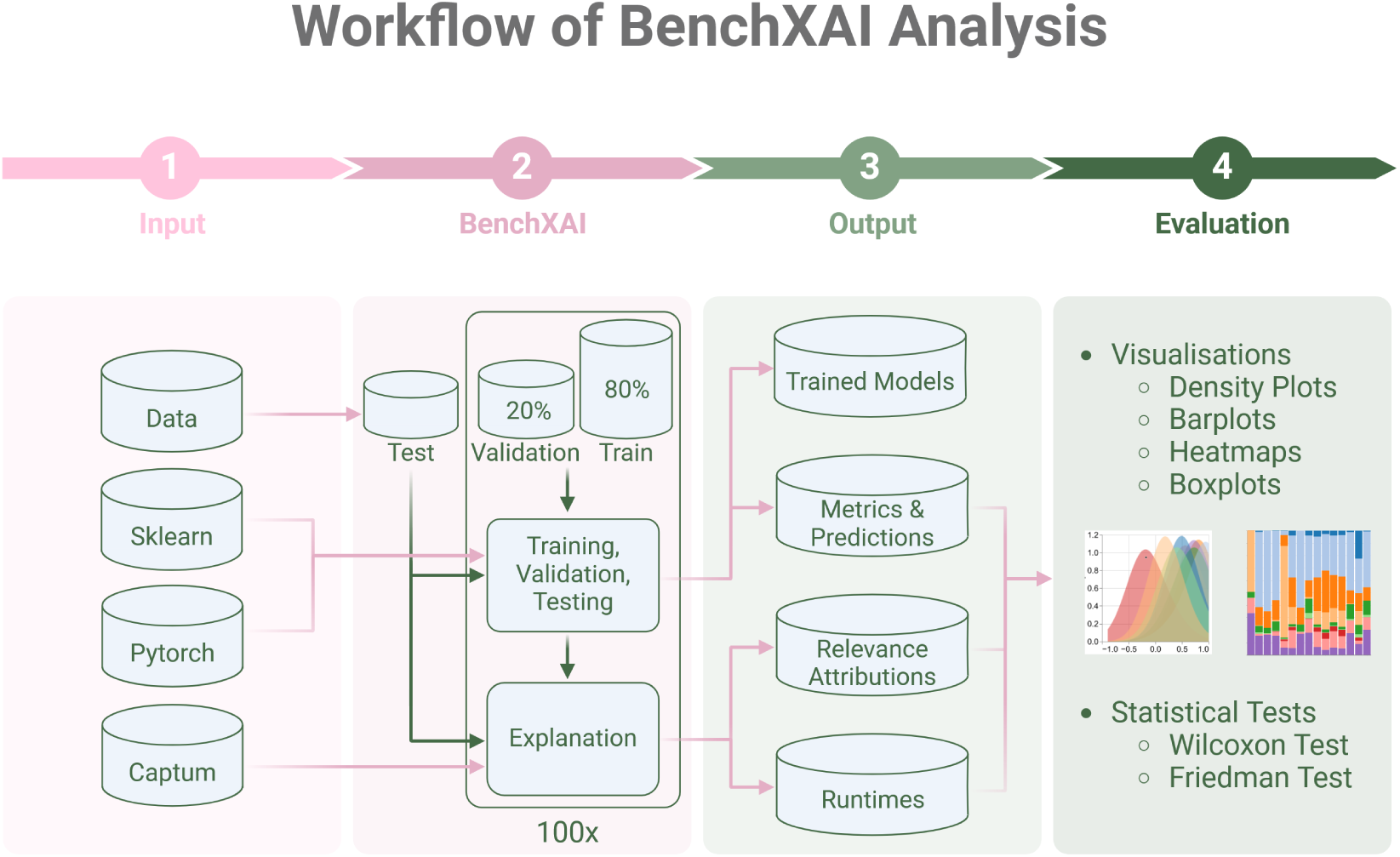
Graphical depiction of the workflow of our BenchXAI framework. In the first step, datasets, machine learning models (ML), deep learning models (DL), and XAI methods are loaded. In the second step our BenchXAI framework is used to split some samples of each class from the dataset for testing and then perform a Monte Carlo Cross Validation (MCCV) followed by the application of 15 different XAI methods on the test data using the trained DL models. In step 3 of our BenchXAI framework all results of step 2 are saved. The fourth and last step is the evaluation of all results.

To help the advancement of segmentation algorithms in ophthalmology, multiple physicians created segmentation masks as part of the **Indian Diabetic Retinopathy Image Dataset (IDRiD)** [39]. The early detection of Diabetic Retinopathy (DR) is crucial in preventing vision loss [1]. A trigger for DR is diabetes, and it is one of the most common causes of preventable blindness [41]. The IDRiD dataset consists of 516 images, but we removed two duplicates resulting in 514 images. 168 images do not show signs of DR, and 346 images show signs of DR. Medical experts graded all images for DR from 0 (no apparent DR) to 4 (severe DR). We combined grades one to four to class 1 (DR) and grade zero was set to class 0 (no DR) to create a binary classification task. The dataset contains pixel-level annotations of abnormalities associated with DR, like microaneurysms, soft and hard exudates, and hemorrhages for 81 images. The images and segmentation masks come in a very high resolution of 4288 × 2848 pixels and were downsized to 858 × 570 pixels using open-cv’s [7] resize function with pixel area relation to decrease runtime and memory usage but still allowing for a good enough DL model performance.

### 2.3 Task 3: XAI Benchmark for Biomolecular Data

Our last task is benchmarking different XAI methods on biomolecular data, such as gene expression data. Biomolecular data is often very high dimensional, leading to issues such as the curse of dimensionality [24] or issues with many highly correlated features. Since there are mostly no gold standards for these data, XAI methods could provide valuable insights in fields such as drug discovery.

The National Institute of Health’s (NIH) National Cancer Institute and the National Human Genome Research Institute brought together **The Cancer Genome Atlas Breast Invasive Carcinoma (BRCA)** data set of breast cancer tissue data from 40 different medical sites. The data consists of annotations about the patient and the tumor alongside raw gene expression counts for each site. We only looked at the raw counts of 20,531 genes for our task and used the molecular sub-type annotation. We combined the data sets from each site into a data set of 1079 samples. To create a binary classification task, our focus will be to predict if a tumor is basal-like (class 0) or luminal-like (class 1), combining the LumA and LumB classes into the luminal-like class and removing the normal and HER2 sub-type classes. We removed samples with missing values, resulting in 867 samples and genes with high correlation (Spearman’s rank correlation coefficient >0.7) and constant features. We used Spearman’s rank correlation because the data is skewed and not normalized. Lastly, we removed all non-coding genes because they only do some regulatory functions as RNA or could be sequencing artifacts, resulting in a total of 11.615 genes. To improve model performance, all genes were normalised using *L*_2_ normalisation. Over 30 gene signatures have been introduced in the last years for breast cancer stratification [21]. The PAM50 assay [36] is a classifier for subtyping breast cancer into luminal, basal-like, normal-like, and HER2-enriched sub-types. In their paper [36], they also show which of the 50 genes are over- or underexpressed in which subtype. Our goal is to verify whether the XAI methods can identify the PAM50 genes that are overexpressed in the luminal subtype.

## 3 Methodology

### 3.1 BenchXAI Package and Workflow

As part of our XAI evaluation, we implemented the BenchXAI framework. It is an easy-to-use framework that allows the user to easily generate XAI attributions for up to 15 XAI methods. We based the framework on Python [57] with Pytorch [37] for Deep Learning (DL) and CaptumAI [26] for XAI attributions, as well as scikit-learn [38] for Machine Learning (ML). The framework currently supports tabular data and image and signal data (see Section 2). The framework comes with an implementation for a Monte Carlo Cross Validation (MCCV) [8], which we use in this study. It is important to note that the **biggest challenge when comparing XAI methods, in particular across different data modalities and datasets, is to achieve the greatest possible comparability, reproducibility, and transferability**. Therefore, we start our analysis by randomly selecting 50 samples of each target class as a test set, which we use to generate relevance attribution. After removing the test set, we randomly split the rest of the dataset into 80% for training the ML and DL models (see Section 3.2) and 20% for validating the models, retaining the distribution of the target classes in each split. To later evaluate how the different XAI methods perform on the models trained on slightly different training sets, we repeat the random 80/20 split 100 times and train the models on each of these 100 splits. This repeated random split is called a Monte Carlo Cross-Validation (MCCV) [8]. Our BenchXAI framework saves all 100 trained models and performance metrics, such as Area under the precision-recall curve (AUPRC), Area under the Receiver Operator Curve (AUCROC), and the Matthews Correlation Coefficient (MCC) for training, validation, and testing, as well as the prediction probabilities for the test samples. The Area under the receiver operating characteristic curve (AUROC) indicates how good the model is at differentiating between the different classes, while the Area under the precision-recall curve (AUPRC) and Matthews Correlation Coefficient (MCC) are better suited for imbalanced datasets [10].

Lastly, we use all 15 XAI methods (see Section 3.3) to calculate attributions for each sample in the test data. This is done for each of the 100 trained AI models, resulting in 100 explanations for each sample in the test set for each XAI method (except for the ShapleyValueSampling XAI method, which had too long runtimes for our image dataset, which is why we were only able to calculate the attributions for 15 iterations). This will allow us to compare the XAI methods against each other and the XAI methods’ performance by themselves on the same test set but with slightly varying underlying AI models. Because the XAI attributions vary strongly, we normalise them before we start our evaluations (see Section 3.4).

### 3.2 AI Models

For our study, we focused on two types of DL architectures for our different classification tasks: the feed forward neural network, also called multi layer perceptron (MLP), and the convolutional neural network (CNN). We have refrained from using recurrent architectures such as Long-Short-Term Memory (LSTM) models since not all of the post-hoc XAI methods presented here can be easily applied to these. Additionally, for Tasks 1 and 3, we trained state-of-the-art ML models, namely random forest and logistic regression models, to compare their model coefficients and feature importance scores to the XAI attributions.

Generally, for all tasks, the DL models had to be specifically tuned to each dataset to achieve a good enough classification performance, which is a crucial baseline for post-hoc XAI methods. If the neural network cannot predict the classes with high enough accuracy, the XAI method might not find reasonable feature importances. For all DL models, the weights are initialized by sampling from a Xavier uniform distribution, and biases are initialized with ones, except for the image and signal data. All models have a linear output layer with 2 neurons for each class. We did not use sigmoid or softmax activations in order to not decrease the network output value, which is used for many XAI methods and might lead to instabilities if too small.

### Task 1: XAI Benchmark for Medical Data

To compare the relevance attributions of the XAI methods to the intrinsically explainable state-of-the-art ML models, we trained a logistic regression and random forest model on the same tabular datasets and extracted their model coefficients for the logistic regression and feature importances (gini impurity) for the random forest. In this study, we used scikit-learns implementation of the logistic regression with default parameters, except for the Heart dataset, where we had to set the regularisation parameter *C* to 100 and the maximum iteration to 1000 in order to achieve better classification results as well as convergence of the algorithm.

The DL model for the WBCD data is an MLP with two hidden linear layers with 20 and 10 neurons, respectively, with ReLU activations, each followed by a dropout layer, with *p* = 0.2. For training, we used cross entropy loss for 300 epochs with a batchsize of 128 and the Adam optimizer with a learn rate of 0.001.

For the Heart data the DL model is a very small MLP with one hidden layer with five neurons and ReLU activation. For training we used cross entropy loss for 1000 epochs with a batchsize of 8 and the Adam optimizer with a learn rate of 0.01.

### Task 2: XAI Benchmark for Medical Image and Signal Data

The DL model for the Signal data is the same as described in Turbé et al. [56]. It is a CNN made up of three blocks, each consisting of a 1D convolutional layer with 256 output channels, a kernel size of 11, a stride length of 1, and a ReLU activation. The three blocks are followed by 1D max pooling with a kernel size of 2, stride length of 2, dilation of 1, and no padding. Lastly, adaptive average pooling and flattening are applied. For training, we used cross entropy loss for 250 epochs with a batchsize of 512 and the NAdam optimizer with a learn rate of 0.00001.

For the IDRiD data, we chose a DenseNet architecture, which is widely used in image classification tasks and was already used for this dataset [40]. We used DenseNet121 with pretrained weights from IMAGENET1K_V1. It has 121 layers, followed by a linear hidden layer with 512 neurons and ReLU activation and a dropout with *p* = 0.1. For training we used cross entropy loss for 200 epochs with a batchsize of 32 and the NAdam optimizer with a learn rate of 0.0001.

### Task 3: XAI Benchmark for Biomolecular Data

Identically to Task 1, we also trained a logistic regression model and random forest model with default parameters for this task. For the BRCA data the model is a small MLP with two hidden layer with 1024 and 128 neurons respectively, with ReLU activations and each followed by a dropout layer with *p* = 0.2. For training, we used cross entropy loss for 5000 epochs with a batchsize of 16 and the Adam optimizer with a learn rate of 0.0000001.

### 3.3 XAI Methods

In this study, we focus on local post-hoc XAI methods. These methods calculate relevance attributions for one specific input sample without considering all other samples in the dataset after the model training. These post-hoc XAI methods are different from global XAI methods, which calculate relevance attributions for each feature based on the whole dataset, as well as ante-hoc XAI methods, which calculate the relevance attributions during the model training (e.g., attention weights in a transformer neural network). Some methods, such as Grad-Cam and Guided Grad-CAM, are not used in this study since they are solely applicable to convolutional neural networks (CNNs) and thus not usable for some of our benchmarking tasks. Throughout this study, we focused on baseline models/values for all XAI methods to validate their usability without extensive hyperparameter tuning. Since post-hoc XAI methods can be very sensitive to these changes [28, 60], data-specific hyper-parameter tuning for each XAI method was not feasible within the scope of this study.

#### 3.3.1 Backpropagation-based methods

Backpropagation-based XAI methods are methods that use the backpropagation mechanism of neural networks to either backpropagate the gradient of the network with respect to the input or some other value, like the network output. Through this, they are able to distribute attributions through the network to the input.

Simonyan et al. [51] introduced the *Input* × *Gradient* method in an image classification specific task to create saliency maps for images as explanations. However, they used the absolute values of the gradients. Shrikumar et al. [49] mentioned this method only incidentally, while demonstrating their DeepLIFT method. However, using the gradient is quite apparent since the gradient tells us how much a change in each input dimension will increase or decrease the output of our model. Multiplying this by the value of the input dimension then gives you an attribution that represents the importance of the input dimension for the output (gradient of input dimension) and how strongly it is represented in the input (value of input dimension). This method is very fast to compute since the gradient is calculated during backpropagation anyway. However, using the gradient is a very naive approach since it might just point to a local optimum of the model output and thus be misleading.

Similar to Input × Gradient, *Deconvolution* [62] computes the gradient of the model output with respect to the input. The methods are identical except that they apply a rectified linear unit (ReLU) activation function to the relevances during backpropagation if a ReLU activation was used in the layer for the forward pass through the network.

*Guided Backpropagation* [52] combines the Input × Gradient [49] method with the Deconvolution [62] method by only backpropagating the relevances when the activation in the forward pass was positive, and the gradient in the backward pass is positive.

Sundararajan et al. [55] introduced the *Integrated Gradients*, which computes the average gradient along a straight line path from a baseline to the given input. Captums [26] default step size of 50 and Gauss-Legendre approximation for the integral was used in this study, as well as, a zero baseline.

Bach et al [4] introduced *Layerwise Relevance Propagation* (LRP) as a method that computes relevance attributions by decomposing the output of a neural network in a backward fashion through all layers of the network until it reaches the input dimension. The decomposition is done in a way that, for any network layer, all relevance attributions for each neuron in that layer always add up to the network output. They propose adding a small stabilising *ɛ >* 0, known as *LRP-ɛ* to prevent unbounded relevance attributions due to possible very small relevances. Captums [26] default LRP method uses the epsilon stabilizer with a small default *ɛ* = 1*e* − 9. To test the influence of the stabilizer we used a larger *ɛ* = 1*e* − 4 for the LRP-*ɛ* method.

Another stabilising decomposition proposed by [4], differentiates between positive and negative pre-activations and only uses the positive pre-activations. This method is known as *LRP-α* − *β*. We used *α* = 1 and *β* = 0, which lays importance only on positive relevances during the backpropagation.

Another decomposition was introduced in [32], which gives more importance for positive relevance attributions, called *LRP-γ*. We use *γ* = 0.25.

Other derivatives of LRP methods, such as LRP-*ω*^2^ [33] are not considered in this study since they are not implemented in the Captum [26] platform, as well as using different rules for different layers in the network [25], since this would require extensive testing of different rule compositions for each network.

*DeepLIFT* (Deep Learning Important FeaTures), introduced by Shrikumar et al. [48], works similarly to the Integrated Gradients method. Integrated Gradients computes the average partial derivative of each feature along a straight line path from input to baseline. At the same time, DeepLIFT approximates this quantity in a single step by replacing the gradient at each non-linearity with its average gradient. We use Captums [26] default zero baseline.

#### 3.3.2 Perturbation-based methods

Perturbation-based XAI methods change an input sample and see how that affects the neural network’s output. One way to do this is to set specific features or amounts of pixels in an image to a baseline value (often zero). In the context of image classification, *Occlusion* was done by Zeiler et al. [62]. For tabular data, this method masks one feature in the input, runs a forward pass through the network, and then computes the difference between the outputs as the relevance attribution. For image data, this can be done for each pixel individually or by selecting a feature mask that slides over the image and sets all pixels underneath the window to a baseline value and then calculating the difference in outputs. We use Captums [26] default zero baseline. For the image data with three color channels, we use a feature mask of shape (3, 3, 3) with step size 3. For the signal data, we use a feature mask of shape (1, 4) with step size 1 in the feature dimension and step size 4 in the time dimension, leading to aggregation only happening along the time dimension.

Shapley values were derived in 1952 by L. S. Shapley [46] and then later applied in the context of cooperative game theory by Shapley and Shubik [45]. Shapley values compute the average marginal contribution of a feature value across all possible feature subsets. To overcome the computational complexity of calculating the Shapley values, Strumbelj et al. [54] introduced a sampling based approach that allows for faster computation of Shapley values, called *Shapley value sampling*. We use Captums [26] default parameters for this method.

Lundberg and Lee [44] proposed SHAP (SHapley Additive exPlanation) values as the Shapley values of a conditional expectation function of the original model. *GradientSHAP* [44] is a method that approximates SHAP values by computing expectations of gradients through random sampling from a distribution of baseline values, which can be seen as an approximation of the Integrated Gradients method. We use Captums [26] default parameters.

*KernelSHAP* [44] uses the LIME method [42] to approximate SHAP values. We use Captums [26] default parameters, except for the signal and image data, where we used a feature mask of (1, 4) and (5, 3), respectively. Another derivation of SHAP is *DeepLiftSHAP* [44], which uses the DeepLIFT method [48] to approximate SHAP values by rewriting the DeepLIFT multipliers in terms of SHAP values (see [44] for the concrete formulas). We use Captums [26] default parameters.

#### 3.3.3 Surrogate methods

Surrogate XAI methods train explainable AI models (e.g., linear regression) to approximate the neural network. Since this is not possible because of the complexity of neural networks, they only approximate it locally around the input sample, allowing for a sufficiently good approximation and interpretability of the surrogate model parameters.

Ribeiro et al. [42] implemented the novel *LIME* (Local Interpretable Model-agnostic Explanations) method, which trains an interpretable model locally around the input prediction. The idea behind LIME is a fidelity-interpretability trade-off, meaning the interpretable surrogate model should be faithful in approximating the black-box model locally while having a low enough complexity to still be interpretable by humans. The method is creating perturbed versions of the input, observing how the model’s predictions change, and subsequently fitting the interpretable model, called surrogate, to these perturbations. LIME is a model agnostic method since it can be applied to any AI method.

We use Captums [26] default values for LIME, namely, scikit-learns [38] Lasso function as an interpretable method with *α* = 0.01, a function which applies an exponential kernel with width 1.0 to the cosine distance between the original input and perturbed input as the proximity measure, and a perturbation function that selects each element independently and uniformly at random. We used a feature mask of (1, 4) and (5, 3) for the signal and image data, respectively.

### 3.4 XAI Evaluation

Since comparability, reproducibility, and transferability of XAI methods across different data modalities and datasets remain one of the biggest challenges when comparing XAI methods, we created a workflow using Monte Carlo Cross Validation (MCCV) to create a standardised test set for the evaluation of XAI methods (see Section 3.1). The focus of our evaluation lies in the results for the correct classified class 1 test samples. The same analyses could be performed for class 0 and for incorrectly classified samples, but this would result in a very different comparison, which was not the main goal of this study. To analyse and compare the XAI attributions and see which features might have been the most important in the prediction of class 1, we used different statistical methods for each benchmarking task. Additionally, we compare the runtimes of all XAI methods on all datasets to analyse how well they scale to different datatypes and sizes. The following evaluations are applied to all XAI methods for easier comparison.

### Normalisation of XAI relevance attributions

When looking at the raw XAI attributions from each XAI method, we found that they vary widely in range between the methods themselves, between the 100 iterations, and between the test samples. Since the XAI methods used in this study are local attribution methods (see Section 3.3) we have decided to normalise the attributions sample wise using a type of rank normalisation that will keep the ordering of the attributions within each patient.

**Definition 1** (Rank normalisation).

*Let* 1 ≤ *n, m* ∈ N*, D* ⊆ R*^n×m^ denote a data set and X* ⊆ *D a set of test samples. For a test sample x* ∈ *X and a fixed XAI method let x_r_* ∈ R*^n×m^ denote the methods specific relevance attributions for sample x. The rank normalized relevance attribution of x is defined as*

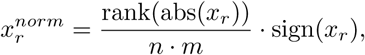

*where the* abs*-function and* sign*-function are applied component-wise and the* rank*-function is defined as follows: Let* 1 ≤ *l* ∈ N*. Given a set of numbers x*_1_*, …, x_l_ with corresponding order statistics x*_(1)_ ≤ · · · ≤ *x*_(_*_l_*_)_ *the subscript* (*i*) *enclosed in parentheses indicates the i-th order statistic of a corresponding number. For all not tied numbers, i denotes their rank, while for all tied numbers the rank is calculated as the arithmetic mean of their corresponding i-th order statistic*.

This way we get relevance attributions in the range of [-1,1] and zero attributions, which indicate no effect, are unchanged. Most importantly, the order of the XAI attributions is not changed through this process, which now allows us to compare the attributions of different XAI methods and between iterations and test samples. *Task 1: XAI Benchmark for Clinical Data*

For the tabular datasets in this task, we looked at stacked density plots to see how the normalised attributions over all correct classified class 1 patients and all 100 iterations are distributed for each feature. Next, we counted how many normalised relevances lie in the range of [−1, −0.5), [−0.5, 0), 0, (0, 0.5], and (0.5, 1] for each feature over all correct classified class 1 test samples and all 100 iterations. We plotted this in a stacked bar plot. These plots show more clearly if some features have over all more positive or negative attributions.

Additionally, we counted how often each feature had the highest positive and lowest negative attribution over all correct classified class 1 test samples and all 100 iterations and plotted the counts in a stacked bar plot. If a feature has the highest attribution very often, that can be a strong indicator that this feature was of great importance in the prediction of class 1.

To compare the XAI method attributions to the logistic regression and random forest models, we plotted the exponentiated model coefficients of the logistic regression model *e^βi^* for all 100 iterations as boxplots. Coefficients greater than 1 indicate a positive effect on the odds for class 1, while a value smaller than 1 indicates a negative effect. In the same way, we plotted the feature importance values of the random forest model for all 100 iterations as boxplots.

To gauge how often the differences in the normalised attributions between the features are significant within an XAI method and iteration, we performed repeated Wilcoxon signed-rank tests [50] for each feature combination for each iteration. A significant p-value indicates the rejection of *H*_0_ : median(Feature 1 - Feature 2) = 0 in favor of *H*_1_ : median(Feature 1 - Feature 2) > 0, indicating that Feature 1 is significantly larger than Feature 2. Lastly, all p-values were corrected using the Bonferroni correction [14], to reduce the likelihood of making a Type-1-error (incorrectly rejecting *H*_0_). The sum of significant p-values (<0.05) for each feature combination over all iterations is then plotted in a heatmap to see how often a normalised attribution for a feature was significantly larger or smaller than another feature’s (maximum of the sum is 100 since we had 100 iterations). Lastly, to determine how stable the XAI methods are between the 100 iterations, we performed a Friedman-Test [17] for each feature. The Friedman test is a non-parametric statistical test used for analysing repeated measures data and is a non-parametric alternative to the repeated measures ANOVA. All p-values were corrected using the Bonferroni correction [14]. Not significant p-values (>0.05) indicate that *H*_0_: „No significant differences in ranked attributions of XAI Method between all 100 iterations.” is not rejected. We show the p-values in a table for each XAI method and each feature and highlight the <u>not significant</u> p-values.

### Task 2: XAI Benchmark for Medical Image and Signal Data

For the image and signal datasets, we plotted the median relevance attributions (median over all 100 iterations) for one random test sample as a heatmap. In the case of the signal data, we plotted the heatmap directly over the signal, while for the image data, we plotted the heatmaps of each XAI method next to the original image. In order to compare the XAI methods over all correct classified class 1 test samples over all 100 iterations, we counted how much each feature of the image/signal is made up of relevance attributions in the ranges of [−1, −0.5), [−0.5, 0), 0, (0, 0.5], and (0.5, 1]. For the IDRiD dataset (see Section 2), this would be the segmentation masks of the different abnormalities, and for the Signal dataset, it would be the different wave patterns (sine, square, and baseline). These are then plotted in a stacked bar plot. Lastly, we plotted the distributions of the relevance attributions of the different features in boxplots for the Signal dataset. For the IDRiD data, we only show the median and interquartile ranges (IQR) values for each feature because there were too many data points to visualise a boxplot.

### Task 3: XAI Benchmark for Biomolecular Data

For the tabular dataset in this task, we first counted how often each feature had the highest positive attribution over all correct classified class 1 test samples and all 100 iterations and plotted the counts in a stacked bar plot (similar to Task 1). Because of the sheer amount of features, this plot more generally shows if some XAI methods behave similarly and not specifically which features might have had the highest importance most often. To get a general overview of features with high or low relevance attributions, we plotted a heatmap of the median normalised attributions for the feature subset defined in our data section, namely the PAM50 geneset [36] (see Section 2) and each XAI method.

To compare the XAI method attributions to the performance of a logistic regression and random forest model (both trained on the same data and with default parameters), we plotted their model coefficients and feature importance scores as boxplots, the same way as for Task 1.

## 4 Results

### 4.1 Task 1: XAI Benchmark for Clinical Data

Overall, the deep learning (DL) and machine learning models (ML) performed quite well according to their area under the receiver operator curve (AUROC), the area under the precision recall curve (AUPRC), and the matthews correlation coefficient (MCC) values, except for on the heart disease data (Heart). The full table with the metric values on training, validation, and test data for all DL and ML models can be found in the Supplementary (see Supplementary Table S3). For the Heart dataset, the neural network was able to better classify class 0 than class 1 (46.94% correct classified class 1 test samples and 90.56% correct classified class 0 test samples, AUROC = 0.743, AUPRC = 0.771, MCC = 0.366). This could be partly due to the small number of overall samples (299) and the imbalance towards class 0 (67,89%) (see Table 1). This struggle of the neural network model to classify class 1 should be considered when looking at the XAI attributions. Lastly, there are only ten class 1 test samples that were correctly classified over all 100 iterations for the Heart data and 29 for the WBCD data.

Figure 4 shows counts on how often each feature has the highest positive attribution for all correct classified class 1 test samples over all 100 iterations. We can see that for the WBCD data (left) BN (light blue) most often has the highest positive attributions for class 1, followed by CT (orange) for most of the XAI methods. Deconvolution and Guided Backpropagation behave differently, giving the highest attribution most often to M (light orange). Additionally, SECS (red) most often has the lowest negative attribution for most XAI methods, meaning it spoke against class 1 the strongest most often (see Supplementary Figure S1). For the Heart data (right), we can see that for most of the XAI methods, Age (light blue) has the largest amount of highest positive attributions, followed by SC (purple) and Sex (brown). Exceptions to this are Deconvolution and Guided Backpropagation, where EF (green) most often has the highest positive attribution. HBP (light green) also has high counts of highest attributions for Input×Gradient, LRP, LRP-*ɛ*, and LRP-*γ*. Only LRP-*γ* additionally has a higher amount of highest attributions for SS (light purple). Lastly, SS (light purple) most often has the lowest negative attribution for most XAI methods, meaning it spoke against class 1 the strongest most often (see Supplementary Figure S1). These results are also reflected when looking at the overall distributions of the XAI relevance attributions for each feature (see Supplementary Figures S2 and S3).

**Figure 4:**
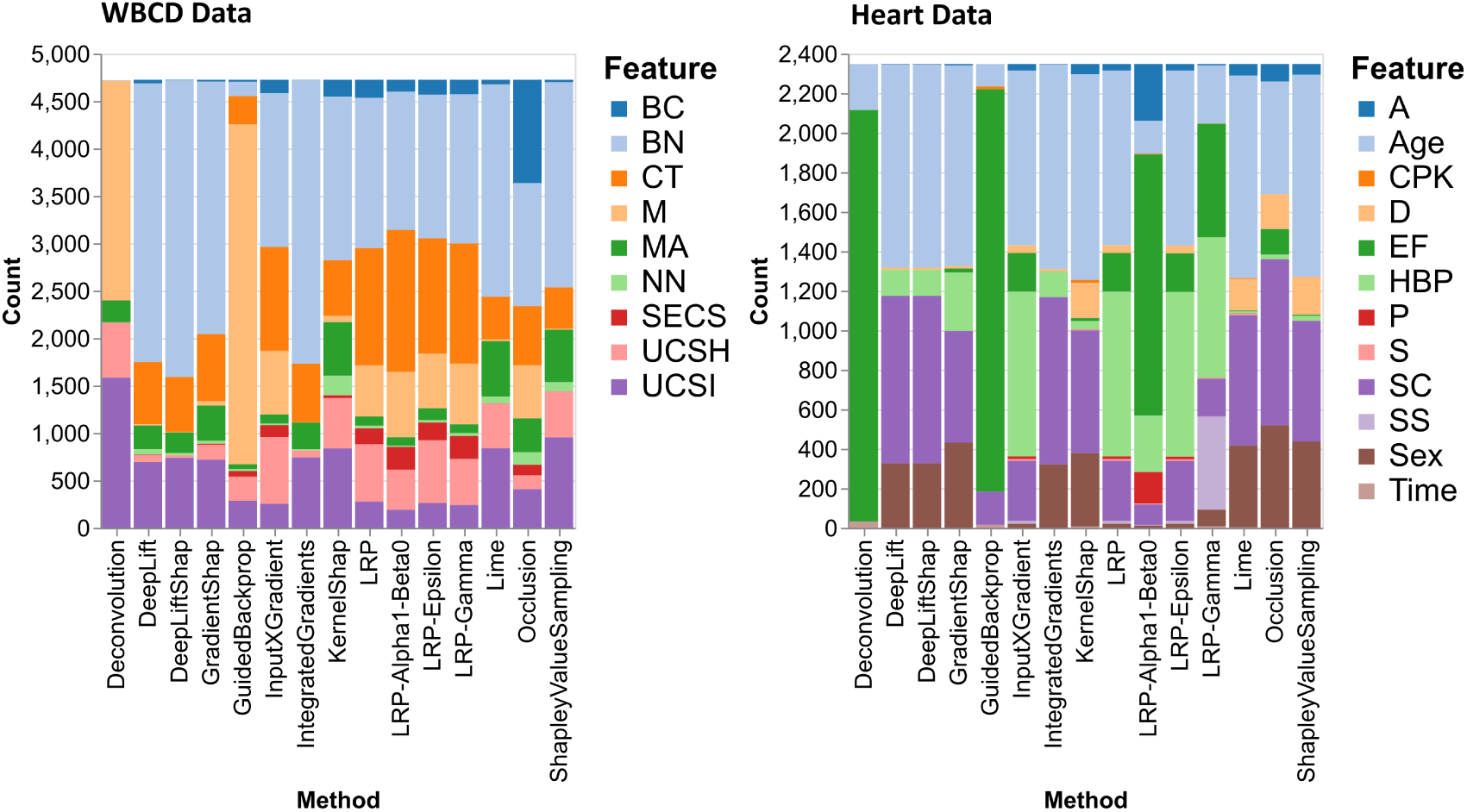
Stacked bar plots for the WBCD dataset (left) and the Heart dataset (right) that show how often each feature had the highest positive attribution for all XAI methods and for all correct classified class 1 test samples over all 100 iterations.

To get an overview if a feature generally has a more positive, negative, or no influence on the prediction, we plotted their counts over all 100 iterations. In Figure 5, we can see that for the WBCD dataset (left), SECS has mostly more negative than positive attributions, followed by M, which also has a few negative attributions. All other features appear to have more positive attributions. LRP-*α*1-*β*0 has the most zero attributions compared to the other XAI methods, and Deconvolution has much more negative attributions for BC, BN, CT, and NN (see Supplementary Figure S4). Overall for the WBCD dataset, Integrated Gradients, DeepLift, GradientShap, and DeepLiftShap perform similarly, as well as Lime, KernelShap, and ShapleyValueSampling, and lastly, LRP, LRP-*γ*, LRP-*ɛ*, GuidedBackpropagation, Occlusion, and Input×Gradient (see Supplementary Figure S4).

**Figure 5:**
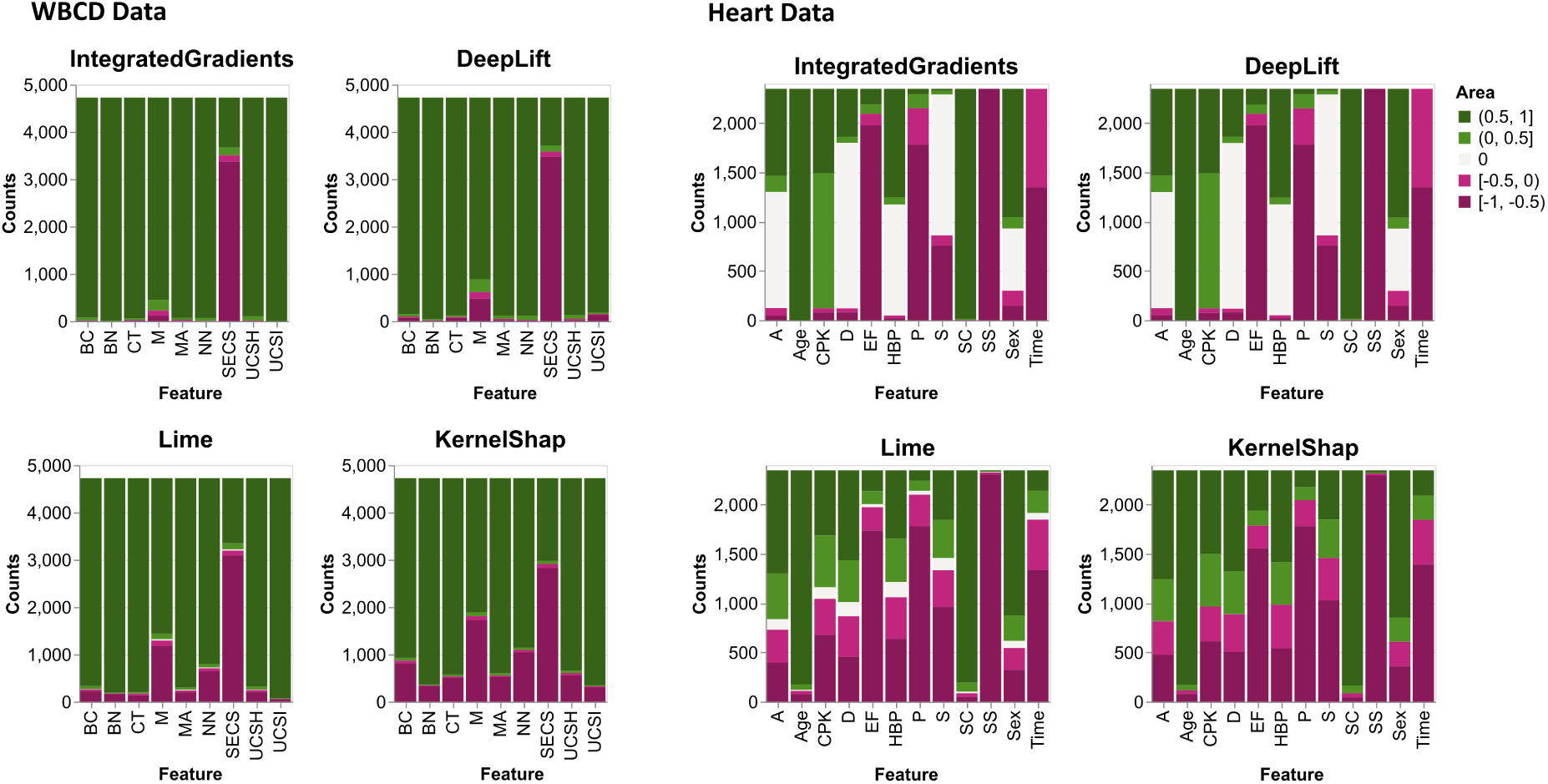
Stacked bar plots for the WBCD dataset (left) and the Heart dataset (right) that show counts of negative, positive and zero attributions for all correct classified class 1 samples and for XAI methods Integrated Gradients, DeepLift, Lime, and KernelShap. All other plots can be found in the supplementary.

For the Heart dataset (right), we can see that for Integrated Gradients and DeepLift, the features A, D, HBP, S, and Sex have a lot of zero attributions, while the SC, Age, and CPK features have overall mostly positive attributions. For Lime and KernelShap, the features Age and SC have mostly positive attributions, and there are fewer zero attributions than for the Integrated Gradients and DeepLift method. LRP-*α*1-*β*0 has more zero attributions, and for Deconvolution and Guided Backpropagation, CPK, EF, and Time have mostly positive attributions (see Supplementary Figure S5). For the Heart dataset, most XAI methods behave similarly to the Integrated Gradients and DeepLift method, except, KernelShap, Lime, LRP-*α*1-*β*0, Deconvolution, and Guided Backpropagation (see Supplementary Figure S5).

To gauge how often the differences in the normalised attributions are significant, we performed repeated Wilcoxon tests for each feature combination and each iteration for all correct classified samples. In Figure 6, we can see that for the WBCD data (left), the attributions for most features are significantly larger than the attributions of SECS, which most often has the lowest negative attribution (see Supplementary Figure S1), and for M. This holds for most XAI methods, except Guided Backpropagation and Deconvolution, as well as, the LRP methods and Input×Gradient, which only has a few or no significant differences (see Supplementary Figure S6).

**Figure 6:**
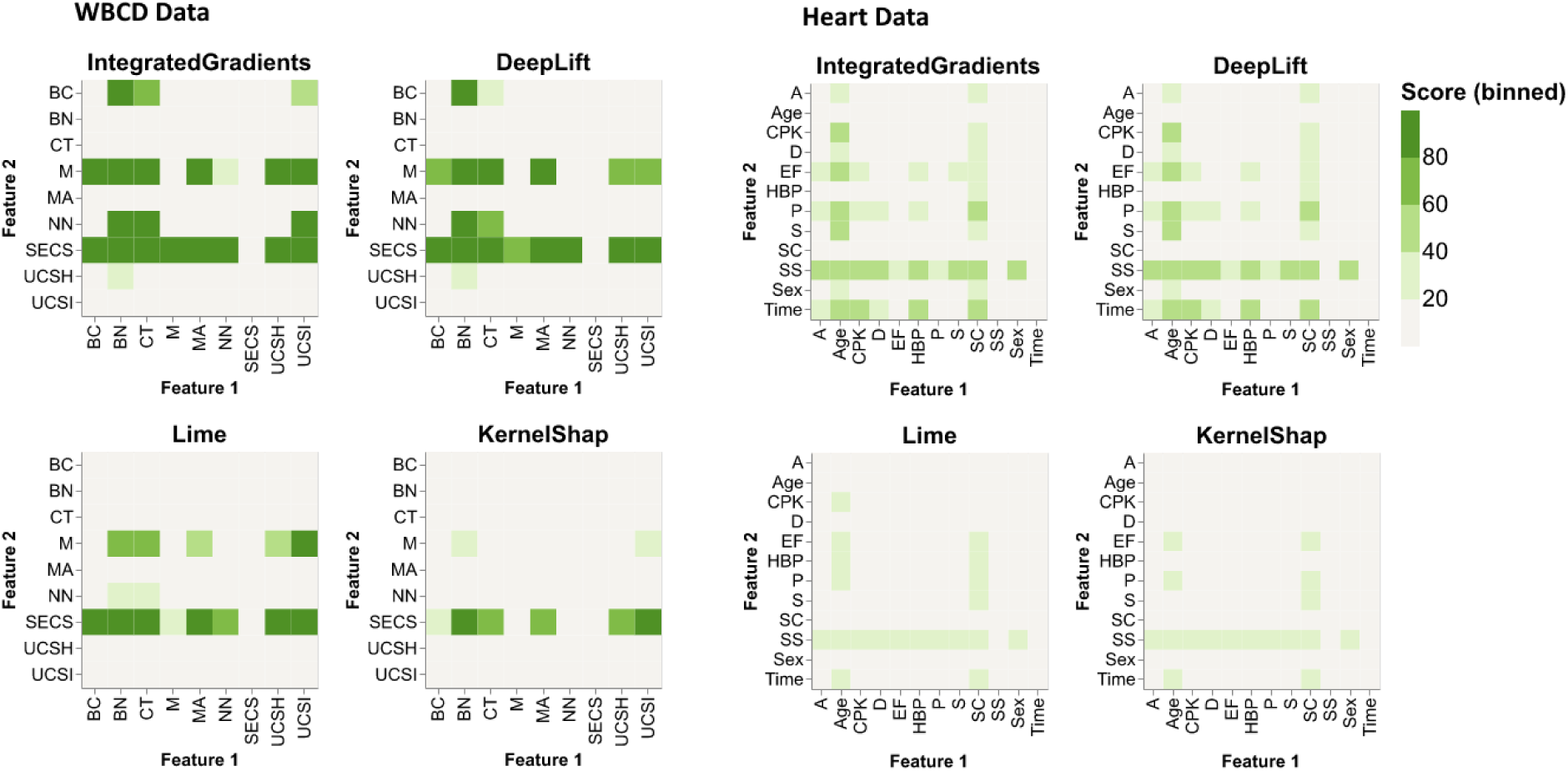
Sum of significant p-values (<0.05) when performing a one sided Wilcoxon test between all features for all 100 iterations for Integrated Gradients, DeepLift, Lime, and KernelShap for the WBCD data (left) and Heart data (right). A significant p-value indicates the rejection of *H*_0_ : median(Feature 1 - Feature 2) = 0 in favor of *H*_1_ : median(Feature 1 - Feature 2) > 0, indicating that feature 1 is significantly larger than feature 2. P-values were corrected using Bonferroni correction to account for multiple testing. All other plots can be found in the supplementary.

For the Heart data (right), we can see that there are fewer significant p-values than for the WBCD dataset (left). For most of the XAI methods, most features are often significantly larger than SS, which has the lowest negative relevance attribution very often (see Supplementary Figure S1). For most XAI methods, except LRP-*α*1-*β*0, Age and SC are sometimes significantly larger than most of the other features (they also most often had the highest positive attribution for most XAI methods (see Figure 4). Similar to the WBCD data, the LRP methods and Input×Gradient only had a few or no significant differences (see Supplementary Figure S7).

Lastly, we wanted to analyse the stability of the XAI methods between the 100 iterations. For this, we performed repeated Friedman tests for each feature. This test determines whether there are significant differences in the ranked relevance attributions of one XAI method for one feature between the 100 iterations. In Table 2, we can see that for the WBCD data for feature CT, there is no significant difference in the normalised attributions between the 100 iterations for all XAI methods except Deconvolution and LRP-*α*1-*β*0. This is followed by BN with no significant differences for most XAI methods, except Deconvolution, Input×Gradient, LRP, and LRP-*ɛ*. Looking at it from the XAI methods, KernelSHAP has no significant differences between the 100 iterations for all features, followed by Occlusion with no significant differences except for features M and SECS. Deconvolution had significant differences for all features. With the results from the stacked barplots (see Figure 4) this could indicate that XAI methods are more stable in the features which most often had the highest importance and more unstable in the other features, whereas KernelSHAP could generally be more stable.

**Table 2:** Bonferroni corrected p-values of repeated Friedman tests for each feature and XAI method. Test were performed on all 29 class 1 samples that were correctly classified over all 100 iterations. All NOT significant p-values (>0.05) are highlighted in green. These indicate that *H*_0_ : „No significant differences in ranked attributions of XAI Method between all 100 Iterations.” is not rejected.

| | BC | BN | CT | M | MA | NN | SECS | UCSH | UCSI | $\Sigma$ |
| --- | --- | --- | --- | --- | --- | --- | --- | --- | --- | --- |
| Deconvolution | 0.0 | 0.0 | 0.0 | 0.0 | 0.0 | 0.0 | 0.0 | 0.0 | 0.0 | 0 |
| DeepLift | **1.0** | **1.0** | **1.0** | 0.0004 | **0.89** | **1.0** | 0.0005 | 0.0 | 0.0 | 5 |
| DeepLiftShap | 0.0 | **1.0** | **1.0** | 0.0 | 0.0004 | **1.0** | 0.0 | 0.0 | 0.0 | 3 |
| GradientShap | 0.0009 | **1.0** | **1.0** | **0.076** | 0.0 | **1.0** | 0.0021 | 0.0 | 0.003 | 4 |
| GuidedBackprop | 0.0 | **1.0** | **1.0** | 0.0 | 0.0 | 0.0001 | **1.0** | 0.0016 | 0.0 | 3 |
| InputXGradient | 0.0016 | 0.0 | **1.0** | 0.0002 | 0.0 | 0.0 | 0.0 | 0.0 | 0.0 | 1 |
| IntegratedGradients | 0.0 | **1.0** | **1.0** | 0.0 | 0.0 | **1.0** | 0.0 | 0.0 | 0.0 | 3 |
| KernelShap | **1.0** | **1.0** | **1.0** | **1.0** | **1.0** | **1.0** | **0.09** | **0.86** | **1.0** | 9 |
| LRP | 0.0026 | 0.0 | **1.0** | 0.0043 | 0.0 | 0.0303 | 0.0001 | **1.0** | 0.0 | 2 |
| LRP-Alpha1-Beta0 | 0.0 | **1.0** | 0.0 | **1.0** | **1.0** | **1.0** | **1.0** | **0.35** | 0.0 | 6 |
| LRP-Epsilon | 0.0 | 0.0002 | **1.0** | 0.0 | 0.0 | 0.0001 | 0.0 | **0.67** | 0.0 | 2 |
| LRP-Gamma | 0.0 | **0.1** | **1.0** | **1.0** | 0.0 | 0.0015 | **0.1** | **1.0** | 0.0 | 5 |
| Lime | **0.16** | **1.0** | **1.0** | **1.0** | **0.09** | **1.0** | 0.0 | 0.0 | 0.002 | 6 |
| Occlusion | **0.07** | **1.0** | **1.0** | 0.0001 | **1.0** | **1.0** | 0.0 | **1.0** | **1.0** | 7 |
| ShapleyValueSampling | 0.0 | **1.0** | **0.3** | **1.0** | **1.0** | **1.0** | 0.0 | 0.0 | 0.0 | 5 |
| $\Sigma$ | 4 | 11 | 13 | 6 | 6 | 9 | 4 | 6 | 2 | 61 |

For the Heart data, there were overall fewer significant p-values (see Supplementary Table S5), but it should be noted that this could also be due to the small sample size used in the test (only 10 class 1 test samples correctly classified over all 100 iterations). D, A, and SC had the least significant differences between iterations for most of the XAI methods, except Deconvolution, Guided Backpropagation, and GradientShap. This is followed by HBP, Time, and S. Deconvolution had significant differences for all features.

Lastly, we wanted to analyse how the XAI relevance attributions compare to the global model coefficients of a logistic regression model, as well as feature importance values from a random forest classifier.

In Figure 7, we can see that for the WBCD data (left), the exponentiated logistic regression model coefficients have mostly median attributions larger than one, meaning they increase the odds for class 1. Only UCSI has a median attribution below one. M (light orange) has the highest median attribution, followed by CT (orange), BN (light blue), and BC (blue). This is similar to most of the XAI methods where BN, CT, and M most often had the highest positive attribution but not BC (see Figure 4). For the random forest classifier, UCSI (purple) has the highest median attribution, followed by UCSH (pink) and BN (light blue). When looking at the counts of highest attributions for class 1 for all XAI methods (see Figure 4), UCSI (purple) and UCSH (pink) have fewer counts of highest positive attributions, then BN (light blue) and CT (orange). For the Heart data, we plotted the exponentiated model coefficients of the logistic regression model on a log scale because of their range. The medians for all coefficients are larger than 1, except for SS, P, EF, and Time. The highest increase in the odds of class 1 is achieved by SC (pink), followed by Age (blue), CPK (orange), and A (light blue). For most of the XAI methods Age (blue) and SC (pink) most often spoke the strongest towards class 1 (see Figure 4). For the random forest model, Time (light brown) has the highest feature importance. This is followed by SC (pink), Age (blue), CPK (orange), EF (green), P (red), and SS (purple), who all have feature importance values close to each other.

**Figure 7:**
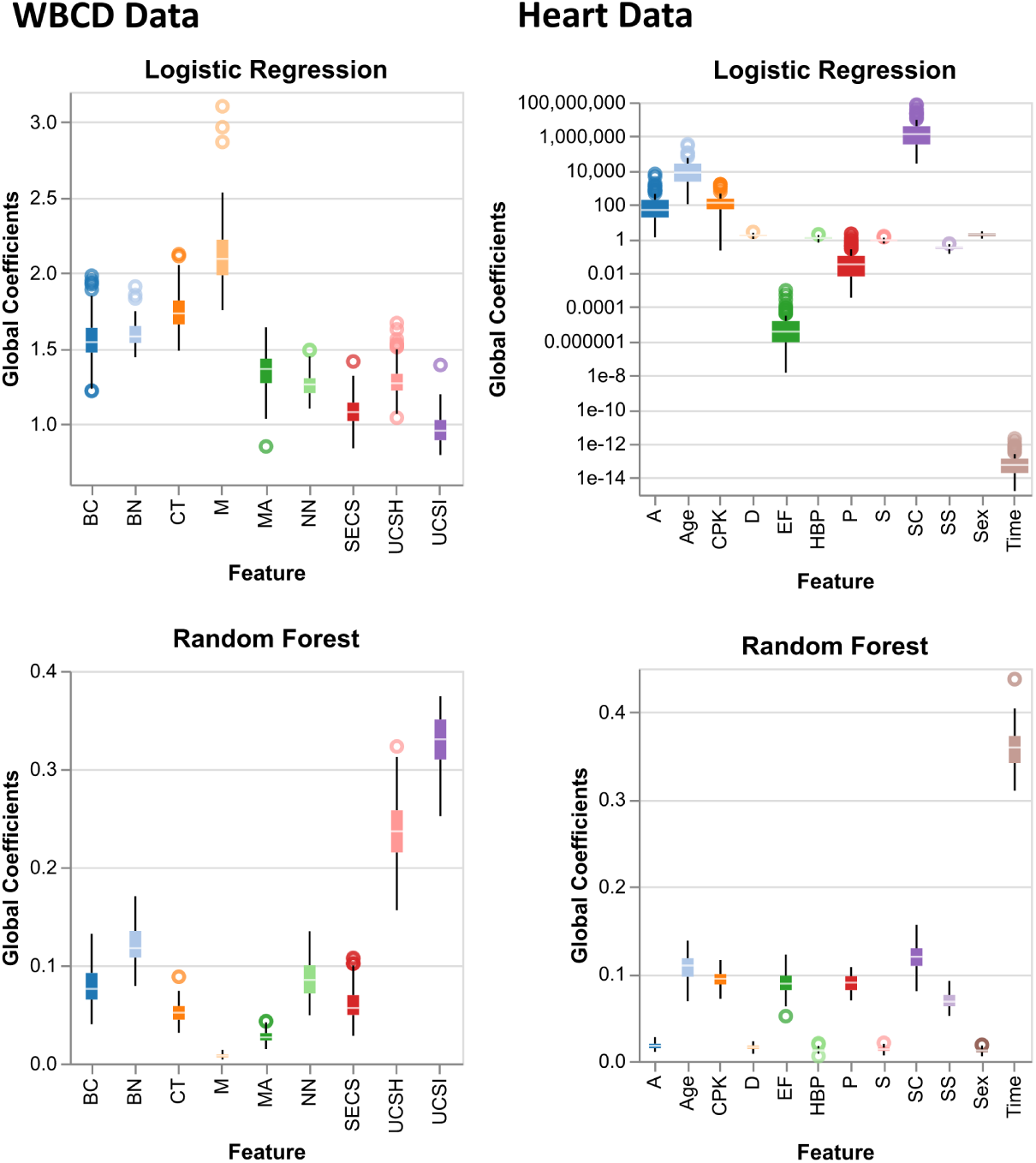
Boxplots of global feature model coefficients and feature importances from logistic regression model and random forest classifier model over all 100 iterations for the WBCD data (left) and the Heart data (right).

In summary, for this task, we showed that most XAI methods behave similarly, except Deconvolution, Guided Backpropagation, and LRP-*α*1-*β*0. For the WBCD dataset, all other XAI methods have mostly low negative attributions for SECS and high positive attributions for BN and CT. BN and CT are also the features found as most important by [18] by their rule extraction algorithm. For the Heart dataset, SC and Age had mostly high positive attributions for all XAI methods except Deconvolution, Guided Backpropagation, and LRP-*α*1-*β*0. For Deconvolution, Guided Backpropagation and LRP-*α*1-*β*0 EF had mostly high positive attributions. Chicco and Jurman [11] identified SC and EF as the most important features, while Ahmad et al. [3] additionally identified Age, HBP, and A as important in the survival prediction.

We showed that the LRP methods and Input×Gradient had very few significant differences in the relevance attributions between the features for this task, while all XAI methods had few significant differences for the Heart data. This is probably due to the fact that the deep learning (DL) model performance for this dataset was not very good (AUROC = 0.738, AUPRC = 0.771, and MCC = 0.366 on the test data), resulting in a smaller sample size of correct classified samples for used for the Wilcoxon test.

Additionally, we showed that most XAI methods have few significant differences between the 100 iterations for the "most significant" features, namely BN and CT for the WBCD data and SC, A, and HBP for the Heart data. It should, however, be noted that the sample sizes for the Friedman test were quite small (29 correct classified samples over all 100 iterations for the WBCD data and 10 for the Heart data), which could influence whether significant differences are found.

Lastly, when comparing the local XAI attributions to the global model coefficients of the logistic regression model, we showed that they perform similarly, except for Deconvolution, Guided Backpropagation, and LRP-*α*1-*β*0. In contrast, the global feature importances of the random forest model varies from the local XAI attributions.

### 4.2 Task 2: XAI Benchmark for Medical Image and Signal Data

All DL models performed exceptionally well on both datasets in this task (AUCROC > 0.98, see Supplementary Table S4 for a full table of evaluation metrics).

For this task, we first looked at the XAI attributions for a random sample of each dataset. For the Signal data (see Figure 8), we can see that Integrated Gradients and DeepLift have mostly positive attributions for the relevant features (Features 3 and 4). In contrast, Guided Backpropagation and Deconvolution have positive attributions for the maxima and negative attributions for the minima of the relevant features. Overall, all XAI methods had positive and negative attributions for the square waves, which were irrelevant to the classification and were added just to confuse the DL model. All other XAI methods behave similarly to the Integrated Gradients and DeepLift methods, except LRP-*α*1-*β*0, which behaves similarly to Deconvolution and Guided Backpropagation. Lime produces overall mostly zero attributions, and KernelShap very small attributions (see Supplementary Figures S12-S14).

**Figure 8:**
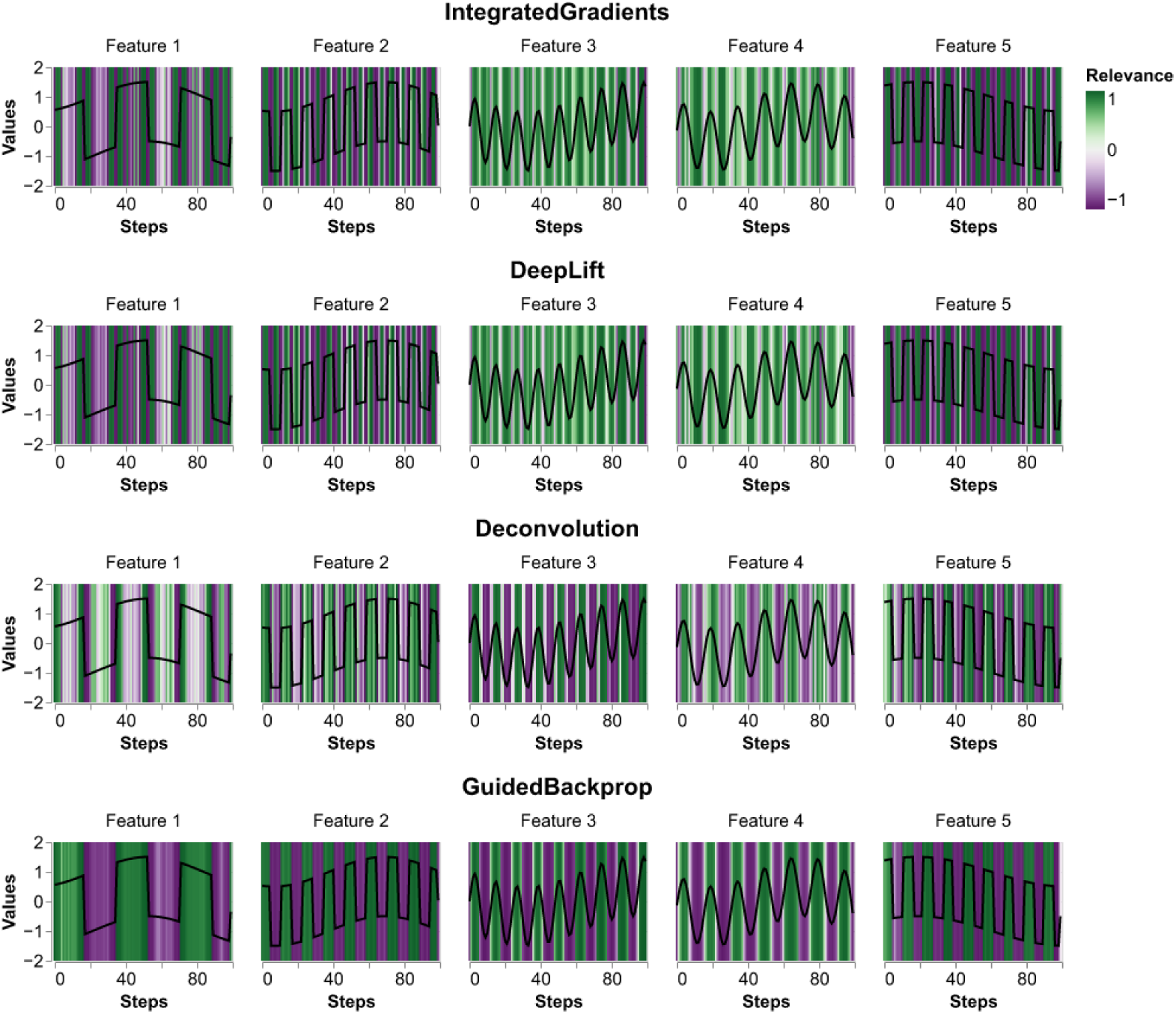
Median relevance attributions over all 100 iterations for one random sample for XAI methods Integrated Gradients, DeepLift, Deconvolution, and Guided Backpropagation. All plots can be found in the Supplementary.

For the IDRiD data (see Figure 9), we can see in the original image the Optic Disc (very bright and on the left side of the eye) as well as Hard Exudates (very bright and more to the middle of the eye). The Optic Disc is not associated with Diabetic Retinopathy (DR), while Hard Exudates (EX) are. Other abnormalities, such as Microaneurysms (MA), we can not easily see. In Figure 9 We can see that all methods were able to identify the shape of the eyeball from the black background, except LRP-*γ* which we did not display, did not converge giving no relevance attributions, while KernelShap and Lime only produced noise or zero attributions over the whole image (see Supplementary Figure S11). Occlusion has mostly negative attributions with some areas, with positive attributions. LRP-*α*1-*β*0 has the most positive attributions, especially around the EX, but also for the OD. It also has a lot of high positive attributions around the edge of the eyeball. Guided Backpropagation, Integrated Gradients, and DeepLift seem to have some negative attributions around the EX but it is not visible if they lie on the EX or right next to it. To further investigate this, we counted how many positive and negative attributions lie within the segmentation masks of the different abnormalities, as well as the sine and square waves of the Signal data.

**Figure 9:**
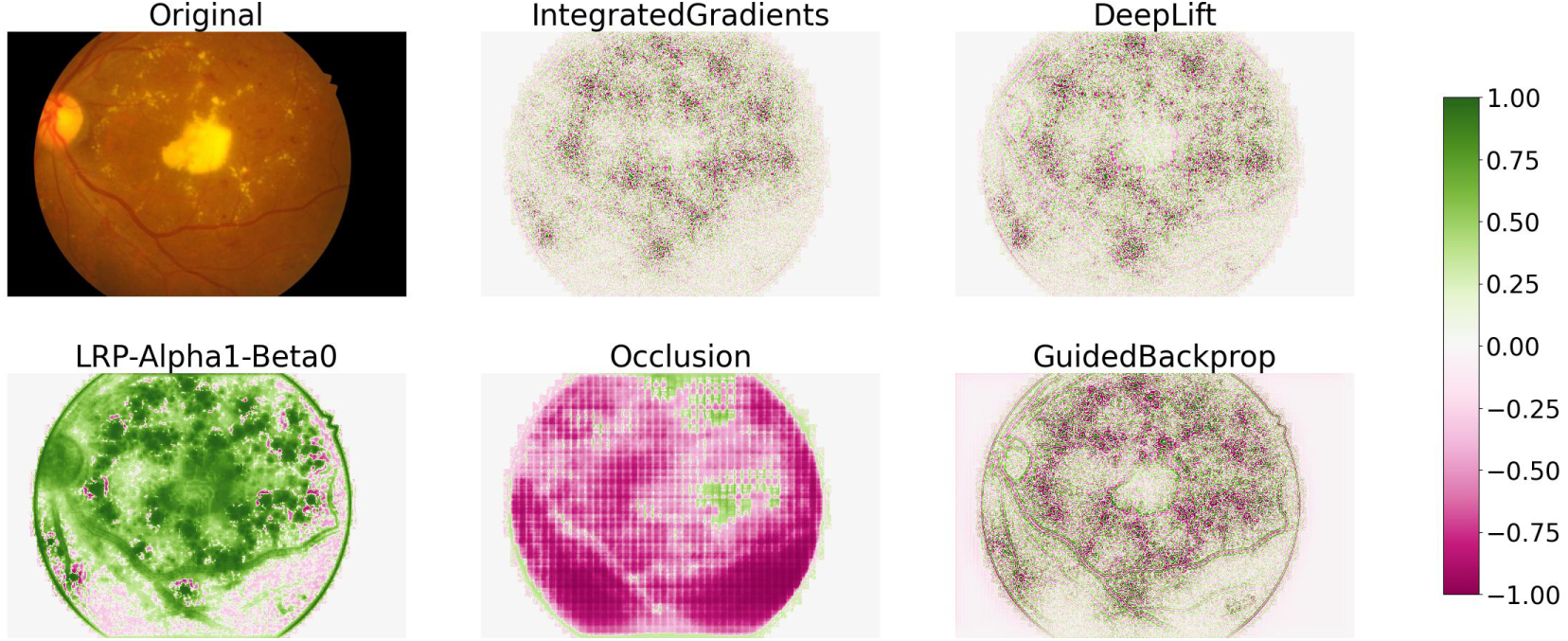
Original image and median relevance attributions for XAI Methods Integrated Gradients, DeepLift, LRP-*α*1-*β*0, Occlusion, and Guided Backpropagation. Medians are taken over all 100 iterations. All plots can be found in the Supplementary.

In Figure 10, we can see that for the Signal data (top), all XAI methods have more smaller positive and negative attributions for the baseline wave and larger positive and negative attributions for the square waves For the IDRiD data (bottom), we can see that for all XAI methods, only the irrelevant parts of the image (Normal) have zero attributions, except Guided Backpropagation (and Deconvolution, see Supplementary Figure S9). These are probably the parts outside of the actual eyeball. For Integrated Gradients and DeepLift, the amount of positive and negative attributions for all abnormalities is around 50%, giving no clear indication whether they were important for the classification of DR. Occlusion has overall more negative attributions for the normal not relevant parts of the images, but also for the abnormalities. LRP-*α*1-*β*0 has more positive attributions for all abnormalities but also for the normal, irrelevant part of the images. Guided Backpropagation has more negative than positive attributions for MA, more positive attributions for EX, OD, and SE, and about the same amount of positive and negative attributions for HE and the normal, not relevant part of the images. All other XAI methods behave similarly to Integrated Gradients and DeepLift, except Deconvolution, KernelShap, and Lime.

**Figure 10:**
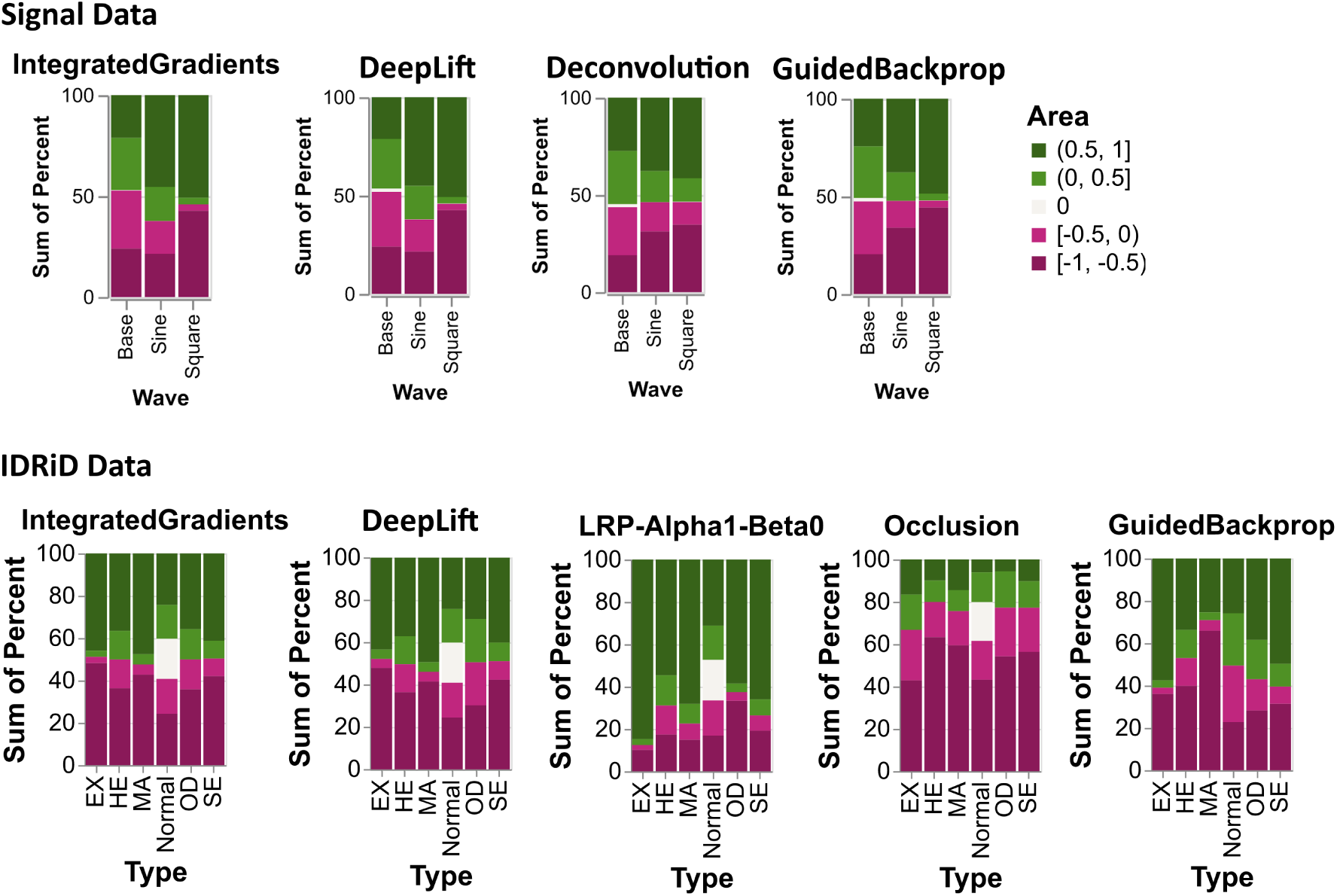
Plots showing how much each abnormality is made up of positive, negative or zero attributions (in percent) for all correct classified class 1 samples. Abnormalities are Microaneurysms (MA), Haemorrhages (HE), Hard Exudates (EX), Soft Exudates (SE), Optic Disc (OD), and Normal for everything outside of the segmentation masks. Plots are shown for Integrated Gradients, DeepLift, Deconvolution, Guided Backpropa-gation, LRP-*α*1-*β*0, and Occlusion XAI methods. All plots can be found in the Supplementary.

Lastly, we wanted to analyse the overall relevance distribution for the features and all XAI methods to see if there may be a trend in the median relevance attribution. For the Signal data, we were able to plot the distributions in the form of boxplots (see Figure 11), but because the images of the IDRiD dataset are very large, we could not plot boxplots. However, we did calculate the median and interquartile ranges (IQR) (see Table 3). In Figure 11, we can see that for all XAI methods, the median attribution for the baseline wave is close to zero with a small interquartile range. For the square wave, the interquartile range is much larger for all XAI methods, and the median is overall more positive, which was to be expected since the square wave had mostly larger positive and negative XAI attributions (see Figure 10). The median for the sine wave attributions is similar to the median of the square wave attributions, but the interquartile range is smaller. The plots for all XAI methods can be found in Supplementary Figure S10.

**Figure 11:**
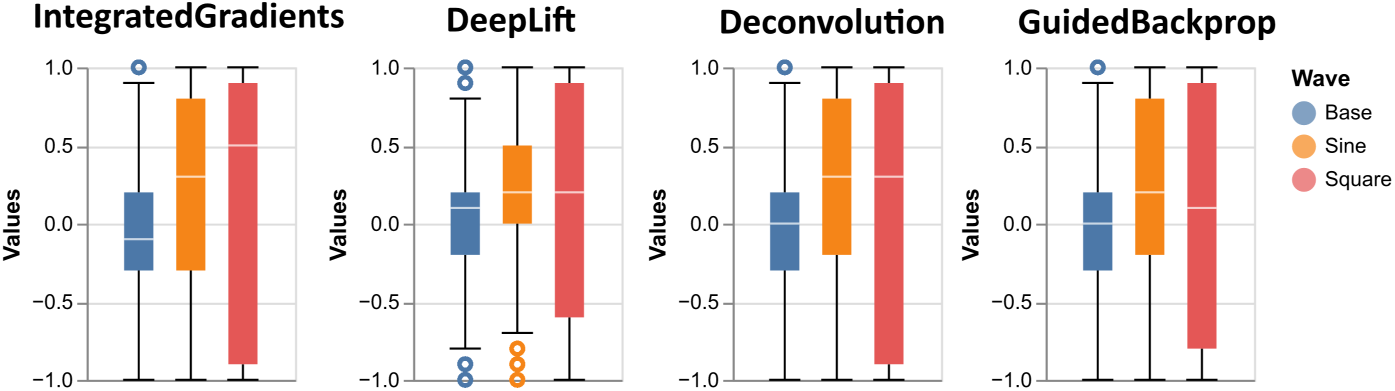
Boxplots of relevance attributions for all correct classified class 1 test samples of the Signal dataset for XAI methods Integrated Gradients, DeepLift, Deconvolution, Guided Backpropagation. All plots can be found in the Supplementary.

**Table 3:** Median and interquartile ranges (IQR) for XAI relevance attributions for all correct classified class 1 samples and all 100 iterations. XAI methods shown are Integrated Gradients, DeepLift, LRP-*α*1-*β*0, Occlusion, and Guided Backpropagation. Abnormalities shown are Microaneurysms (MA), Haemorrhages (HE), Hard Exudates (EX), Soft Exudates (SE), Optic Disc (OD), and Normal for everything outside of the segmentation masks. LRP-*γ* was removed from the table, because it did not converge for this dataset. Lime was removed from this table, because median and IQR were 0 for everything. The complete list for all other XAI methods can be found in the Supplementary.

|  | MA | HE | EX | SE | OD | Normal |
| --- | --- | --- | --- | --- | --- | --- |
| IntegratedGradients | 0.369<br>$\pm 1.369$ | -0.0<br>$\pm 0.926$ | -0.339<br>$\pm 1.386$ | -0.015<br>$\pm 0.97$ | 0.007<br>$\pm 0.81$ | 0.0<br>$\pm 0.7$ |
| DeepLift | 0.429<br>$\pm 1.406$ | 0.005<br>$\pm 0.955$ | -0.365<br>$\pm 1.318$ | -0.303<br>$\pm 0.986$ | -0.029<br>$\pm 0.756$ | 0.0<br>$\pm 0.713$ |
| LRP-$\alpha 1-\beta 0$ | 0.732<br>$\pm 0.423$ | 0.565<br>$\pm 0.484$ | 0.915<br>$\pm 0.197$ | 0.775<br>$\pm 0.433$ | 0.681<br>$\pm 0.263$ | 0.107<br>$\pm 0.569$ |
| Occlusion | -0.582<br>$\pm 0.29$ | -0.615<br>$\pm 0.295$ | -0.451<br>$\pm 0.21$ | -0.557<br>$\pm 0.294$ | -0.54<br>$\pm 0.197$ | -0.48<br>$\pm 0.709$ |
| GuidedBackprop | -0.751<br>$\pm 1.421$ | -0.272<br>$\pm 1.331$ | 0.709<br>$\pm 1.722$ | 0.469<br>$\pm 1.325$ | 0.369<br>$\pm 1.09$ | -0.025<br>$\pm 0.886$ |

In Table 3, we can see that for the Normal feature (everything outside of the segmentation masks), the medians are zero or close to zero for all XAI methods except for LRP-*α*1-*β*0, Occlusion and KernelShap. Similarly, the Optic Disc (OD), which is not relevant for the classification of DR, has medians close to zero for all XAI methods except for LRP-*α*1-*β*0, Guided Backpropagation, ShapleyValueSampling, Occlusion, and KernelShap (see Supplementary Table S6). Microaneurysms (MA) overall have the most high positive median attributions (>0.3) for all XAI methods except for Guided Backpropagation, Occlusion, Deconvolution, ShapleyValueSampling, and KernelShap. Haemorrhages (HE) have median attributions close to zero for most XAI methods except for LRP-*α*1-*β*0, Guided Backpropagation, Occlusion, Deconvolution, and KernelShap. Hard Exudates (EX) have mostly negative median attributions for all XAI methods except for Guided Backpropagation, LRP-*α*1-*β*0, and Deconvolution. For Soft Exudates (SE), median attributions are negative for all XAI methods except Guided Backpropagation, Deconvolution, and LRP-*α*1-*β*0. Overall, Input×Gradient, GradientShap, IntegratedGradients, LRP-*ɛ*, and LRP have similar median attributions, as well as, Deconvolution and Guided Backpropagation, KernelShap and Occlusion, and lastly DeepLift and DeepLiftShap (see Supplementary Table S6).

Lastly, we also looked at the median and IQR for the different severity grades of DR, but there were no notable differences compared to our analysis on all severity grades combined.

To summarize, We found that Captums [26] default Lime implementation produces mostly zero attributions over for the datasets in this task, and LRP-*γ* did not converge with standard parameters for the IDRiD dataset. For both datasets, all other XAI methods have positive and negative attributions in the relevant and not relevant parts of the sample. LRP-*α*1-*β*0 has more positive than negative attributions.

Deconvolution, ShapleyValueSampling, and KernelShap produced mostly noisy heatmaps for the IDRiD dataset. Deconvolution has already been shown to produce noisier heatmaps [43] compared to some other XAI methods. All other XAI methods identified some patterns in the image. Microaneurysms (MA) overall have the highest attributions. According to [59], MAs are present for all severity grades, while the other abnormalities only start appearing from severity grade 2. We found that none of the XAI methods were able to give more positive attributions to the feature masks than the non relevant parts of the image. One reason for this could be due to the fact that we had to downsize the image as well as the segmentation mask, thus losing some information. Another reason could be that the segmentation masks are not exactly the locations that the models are using for their predictions. When looking at Figure 9 we can clearly see that Integrated Gradients,

LRP-*α*1-*β*0, and Guided Backpropagation found some pattern in the image, but that pattern might just not perfectly overlap with the segmentation masks.

For the Signal data, all XAI methods except Guided Backpropagation and Deconvolution have more positive attributions for the relevant parts of the signal. Guided Backpropagation and Deconvolution seem to have identified the pattern of switching between local minima and maxima and, thus, constantly switching between high positive and high negative attributions.

### 4.3 Task 3: XAI Benchmark for Biomolecular Data

The deep learning (DL) and machine learning (ML) models performed very well on the classification task, though the logistic regression and random forest models outperformed the neural network in terms of matthews correlation coefficient (neural network MCC = 0.718, logistic regression MCC = 0.942, random forest MCC = 0.961, see Supplementary Table S3 for a full list of evaluation metrics).

Firstly, to get an overview on if the XAI methods found only a few genes important in the classification of class 1 or if many genes were found important, we looked at how often each gene had the highest positive attribution, just like we did for the datasets in Task 1. In Figure 12, we can see that for most of the XAI methods, there are >70 genes that had the highest positive attribution at least once. KernelShap, Lime, Occlusion, and ShapleyValueSampling have much more genes that had the highest positive attribution at least once. In contrast, Deconvolution and Guided Backpropagation have far fewer genes with higher counts.

**Figure 12:**
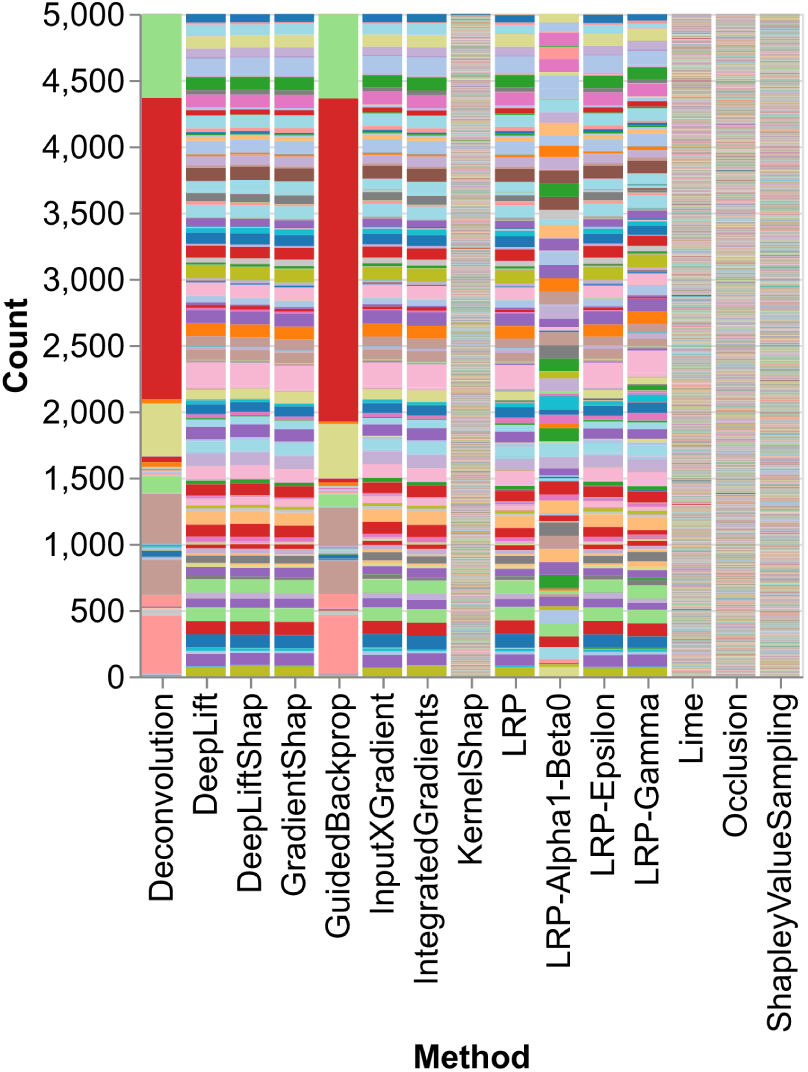
TCGA BRCA data: Stacked barplot that shows how often a feature had the highest positive attribution for all XAI methods, for correct classified class 1 test samples. The legend has purposely been removed, because there are to many features displayed and the colours are repeated, so a comparison of colors is not possible.

Secondly, we wanted to see how relevant the PAM50 genes, which are used for breast cancer subtyping, are for the luminal class (class 1), for all XAI methods. In Figure 13 we can see that KernelShap, Lime, Occlusion and ShapleyValueSampling have relevance attributions close to zero for all PAM50 genes, while LRP-*α*1-*β*0 has positive attributions for all PAM50 genes. All other XAI methods have high median positive attributions for the BAG1, BCL2, BLVRA, CXXC5, ERBB2, ESR1, FOXA1, GPR160, GRB7, MAPT, MDM2, MLPH, MMP11, NAT1, PGR, SLC39A6, and TMEM45B genes and negative median attributions for the other genes. Most of the genes with positive median attributions are overexpressed in the luminal A and/or luminal B subtypes except for BCL2, MDM2, and MLPH, which are overexpressed in the normal-like subtype, and ERBB2, GRB7, and TMEM45B which are overexpressed in the HER2 subtype. We found some genes with negative median attributions that are overexpressed in the luminal B subtype but at the same time underexpressed in the luminal A subtype, namely, UBE2C, PTTG1, MYBL2, CCNB1, TYMS, MELK, CEP55, UBE2T, CDC6, and NUF2 (called CDCA1 in the PAM50 gene set). This could be due to the fact that we combined the luminal A and luminal B subtypes into a luminal-like class for our classification task. All the other genes with negative median attributions are underexpressed in the luminal A and/or luminal B subtypes.

**Figure 13:**
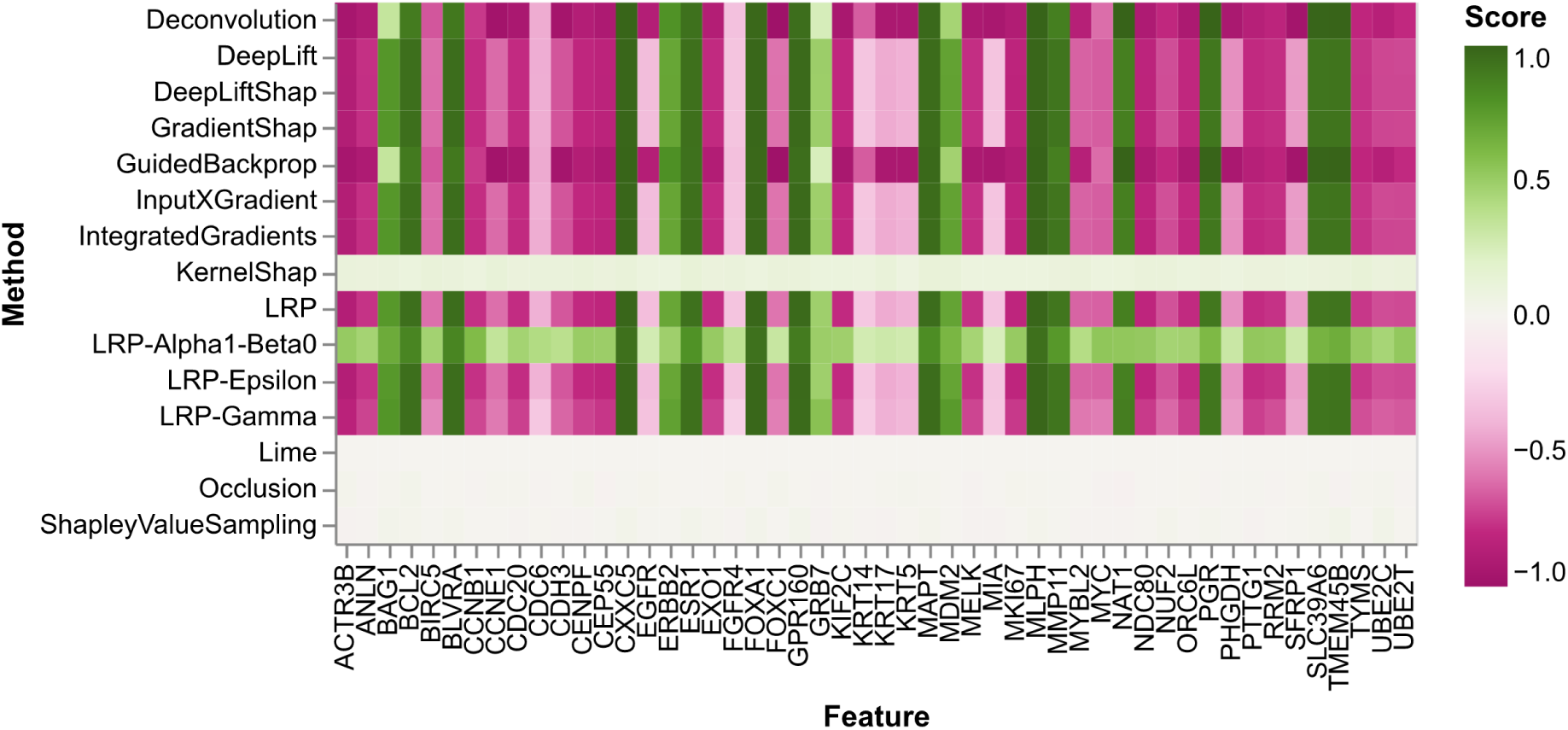
TCGA BRCA data: Heatmap of median attributions for each PAM50 gene and each XAI method. Median is taken over all correctly classified class 1 samples over all iterations.

Lastly, we wanted to analyse how the XAI relevance attributions compare to the global model coefficients of a logistic regression model, as well as feature importance values from a random forest classifier.

For the TCGA BRCA dataset (see Figure 14), the logistic regression model has 18 median model coefficients for the PAM50 genes, which are larger than 1, meaning they increase the odds of class 1. Most of these 18 coefficients also have high relevances for most of the XAI methods, except CDC6, which has low relevances for most XAI methods (see Figure 13), but is actually overexpressed in the luminal B subtype and underexpressed in the luminal A subtype. All of the other 17 model coefficients are overexpressed in the luminal A and/or B subtype except MLPH, BCL2, and MDM2, which are overexpressed in the normal subtype, and TMEM45B, ERBB2, and GRB7, which are overexpressed in the HER2 subtypes. For the random forest model, all median feature importances for the PAM50 genes are small (<0.03). The largest median feature importances (>0.005) are for the ESR1, FOXA1, FOXC1, CXXC5, MLPH, GPR160, and PGR genes of which all had high relevance attributions for most XAI methods, except FOXC1 (see Figure 13). MLPH is not overexpressed in the luminal subtypes but in the normal subtype.

**Figure 14:**
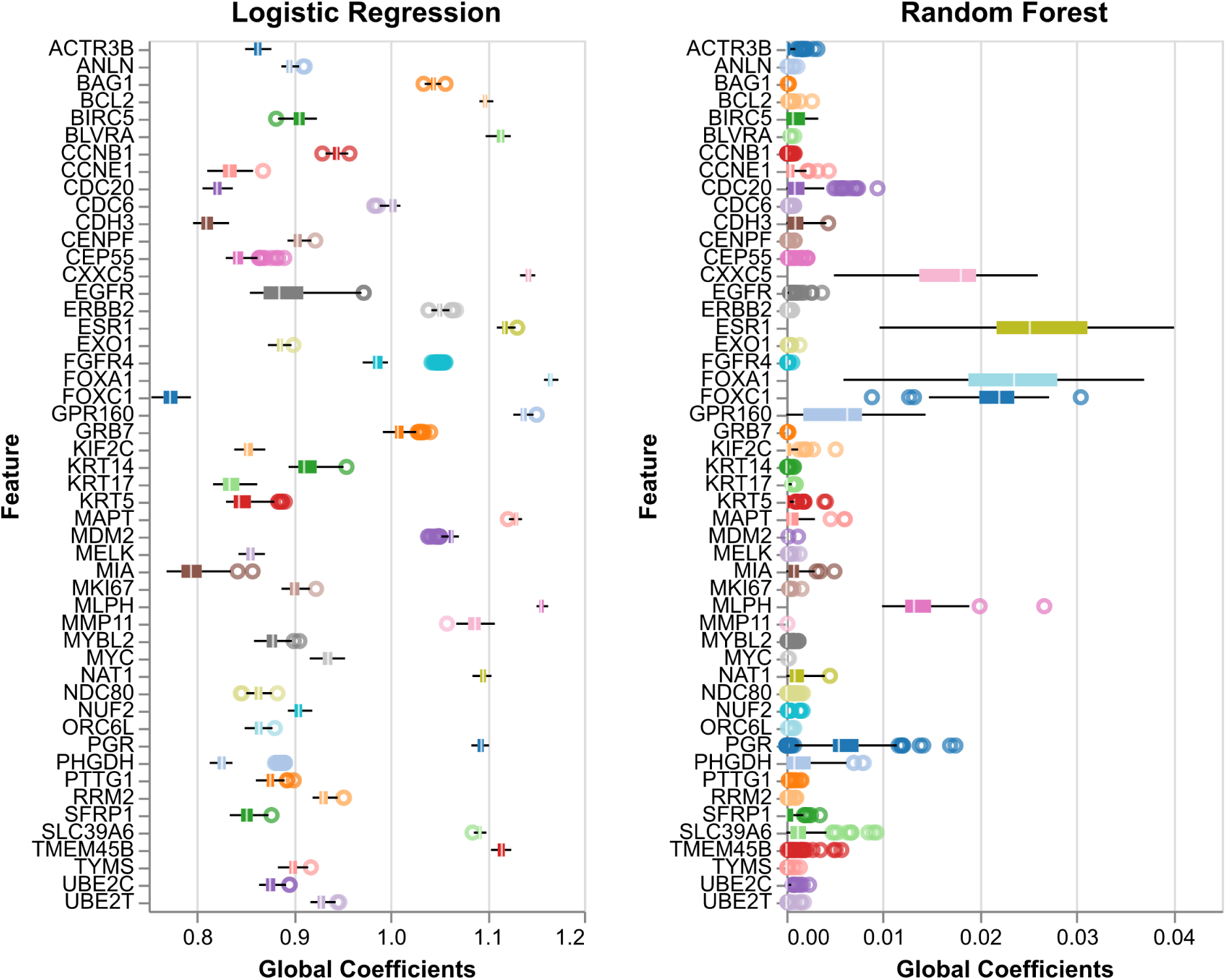
TCGA BRCA data: Boxplots of global feature importances from logistic regression model and random forest classifier model over all 100 iterations for PAM50 genes.

In summary, similarly to Task 1, Deconvolution, GuidedBackpropagation, and LRP-*α*1-*β*0 behave differently from the other XAI methods, as well as KernelShap, Lime, Occlusion, and ShapleyValueSampling for which many genes had the highest positive attribution at least once. This was also reflected in the median attributions for the PAM50 genes, where these four methods had median attributions of close to zero for all PAM50 genes, while LRP-*α*1-*β*0 produced only positive median attributions. All other XAI methods were able to correctly identify BAG1, BLVRA, CXXC5, ESR1, FOXA1, GPR160, MAPT, MMP11, NAT1, PGR, and SLC39A6 which are overexpressed in the luminal A and/or luminal B subtypes [36]. The other six genes with high positive attributions are actually overexpressed in the Normal or HER2 subtypes.

Similarly to Task 1, the global model coefficients of the logistic regression model behave similarly to the XAI attributions, while the random forest feature importance scores do not.

### 4.4 XAI Runtimes

In general, we found that all the runtimes for XAI methods vary depending on the analysed taks. Overall the perturbation-based and surrogate XAI methods (see Section 3.3) had longer runtimes on larger input samples. In Table 4, we can see that ShapleyValueSampling scales very badly to large datasets, with a median runtime of around 2.7 hours for one image in the retina dataset, 1.5 minutes for the BRCA dataset, and 9 seconds for the synthetic dataset. Occlusion had a median runtime of around 10.8 minutes on the retina dataset and 3.4 seconds for the BRCA dataset. KernelShap and Lime had median runtimes of around 5.7 seconds for the retina dataset. All other XAI methods scale very well to larger datasets with median runtimes of less than one second. For the smaller datasets (WBCD and Heart) runtimes were often less than 1 millisecond.

**Table 4:** Median runtimes over all 100 iterations on 100 test samples of all XAI methods for all datasets in milliseconds. All calculations were performed on one NVIDIA RTX5000 GPU. *ShapleyValueSampling runtimes for the Retina dataset were calculated on one NVIDIA A100 GPU and only for 15 iterations due to excessively long runtimes and limited access to the GPU.

| XAI | Task 1 |  | Task 2 |  | Task 3 |
| --- | --- | --- | --- | --- | --- |
|  | WBCD | Heart | IDRiD | Synthetic | BRCA |
| Deconvolution | 0.24 | 0.17 | 34.05 | 0.89 | 0.56 |
| DeepLiftShap | 0.74 | 0.62 | 50.52 | 2.08 | 1.09 |
| DeepLift | 0.46 | 0.34 | 46.85 | 1.89 | 0.97 |
| GradientShap | 0.65 | 0.6 | 30.44 | 1.33 | 0.91 |
| GuidedBackprop | 0.25 | 0.18 | 34.06 | 0.89 | 0.56 |
| InputXGradient | 0.19 | 0.15 | 26.8 | 0.73 | 0.44 |
| IntegratedGradients | 2.37 | 2.25 | 574.46 | 8.19 | 4.21 |
| KernelShap | 68.31 | 59.59 | 5712.39 | 83.38 | 441.13 |
| Lime | 82.28 | 76.49 | 5751.57 | 115.04 | 908.93 |
| LRP-Alpha1-Beta0 | 0.59 | 0.37 | 68.12 | 2.22 | 1.17 |
| LRP-Epsilon | 0.56 | 0.36 | 64.8 | 2.15 | 1.12 |
| LRP-Gamma | 0.62 | 0.38 | 71.4 | 2.29 | 1.21 |
| LRP | 0.56 | 0.37 | 64.88 | 2.17 | 1.13 |
| Occlusion | 1.41 | 1.44 | 648997.59 | 374.64 | 3412.65 |
| ShapleyValueSampling | 46.01 | 52.54 | 9744782.34* | 9090.13 | 92299.58 |
| Data shape | (9) | (12) | (570, 858) | (6, 500) | (867) |

## 5 Conclusion

Comparing different XAI methods on a single dataset is very difficult due to the huge variance in explanations across XAI methods. This is even more true when widening the comparison across different data sets and data modalities. In particular, the inconsistent explanations between XAI methods could potentially reduce trust in AI systems, highlighting the necessity of a large benchmarking study demonstrating the strengths and weaknesses of XAI methods for different applications.

In our study, we aim to address this unsolved issue of evaluating and comparing XAI results across multiple modalities specifically for XAI in medicine.

Therefore, we first developed a targeted benchmarking framework, called BenchXAI, that allows for a comprehensive evaluation of fifteen different XAI methods, focusing on investigating their robustness, suitability, and limitations for various tasks. To achieve this, BenchXAI establishes a generalized evaluation scheme with its own implementation of a Monte Carlo Cross Validation (MCCV), allowing for repeated testing of XAI methods on slightly different AI models due to training on different data splits. Moreover, to investigate the robustness of the XAI methods, BenchXAI implements a multitude of visualisations, such as density plots, barplots, heatmaps, and boxplots, as well as statistical tests, such as the Wilcoxon test and Friedman test. This extensive quantitative analysis enables BenchXAI to thoroughly investigate and compare the performance and robustness of the XAI methods, allowing for a comparison across all ML, XAI, and data modalities.

Most importantly, we introduced a novel sample-wise normalisation method for post-hoc XAI methods (see Definition 1), which maintains the ranking of positive and negative attributions, but scales everything into the range of [−1, 1] leaves the zero attributions untouched so that they can still be interpreted as having no positive or negative effect on the prediction.

Our BenchXAI package thus improves upon current XAI benchmarking packages such as OpenXAI [2] or Xplique [16], which focus only on one data modality (tabular or image data) and evaluate model robustness using only evaluation metrics, for which there currently exists no consensus on which metrics should be employed [22, 19].

Finally, we performed a benchmark comprising a thorough analysis and evaluation of task-specific advantages and disadvantages in three typical biomedical modalities: clinical data, medical image and signal data, and molecular data. We showed that all XAI methods scale quite well to larger data types, except for perturbation-based XAI methods, such as ShapleyValueSampling and Occlusion, which have very large runtimes for bigger input samples, namely image or signal data, which is why we do not recommend their usage for larger data types. Furthermore, we found that XAI attributions of most XAI methods behave similarly to the global model coefficients of a logistic regression model. Overall, we found that XAI methods DeepLift, DeepLiftShap, GradientShap, and Integrated Gradients performed well in all three tasks.

Additionally, based on our results for Task 1, we recommend XAI methods KernelShap, Lime, Occlusion, and ShapleyValueSampling for this task. They were able to find important features that were previously identified by other publications with significantly higher importance than other features and with significant stability over all 100 iterations. We found that the LRP methods give similar relevance attributions to these XAI methods (except LRP-*α*1-*β*0), though with fewer significant differences between relevance attributions of features. As stated in [25], to achieve better results from the LRP methods, they could be applied to different types of layers in a neural network. We found that for clinical data, the XAI methods Deconvolution, Guided Backpropagation, and LRP-*α*1-*β*0 behave differently from all other XAI methods, which is why we do not recommend using them for these tasks, as of yet. Further studies are needed to analyse their relevance attributions.

For signal data, we additionally recommend using XAI methods Input×Gradient, Occlusion, Guided Back-propagation, LRP, LRP-*γ*, and LRP-*ɛ*, though a combination of the LRP methods could lead to better results [25]. These methods had the highest median and lowest interquartile ranges of positive relevance attributions for the relevant parts of the signal. Lime did not work well with default parameters for this type of data. We found that for our dataset, where the relevant pattern was longer (100 time steps), all XAI methods showed positive and negative attributions in these patterns. This makes an automated evaluation almost impossible without looking at the heatmaps for each sample. For this reason, we recommend trying to identify smaller, more localised relevant patterns for a specific task and dataset instead of a larger region.

For medical image data, we additionally recommend using XAI methods Guided Backpropagation, Input×Gradient, LRP, and LRP-*ɛ*, because they were able to identify the relevant patterns of the image, though they did not find many high positive attributions in the actual feature masks. Lime did not work well with default parameters for this type of data, and LRP-*γ* did not converge with default parameters. KernelShap, Deconvolution, and ShapleyValueSampling produced mostly noise for this data. Since none of the XAI methods were able to give more positive attributions to the relevant parts than the non-relevant parts of the image, we recommend performing further studies on segmentation masks for XAI evaluation, such as investigating whether the XAI relevance attributions lie just on the edge of the segmentation masks.

Lastly, for the biomolecular data, we additionally recommend using XAI methods Input×Gradient, LRP, LRP-*ɛ*, and LRP-*γ* because they were able to identify most of the relevant PAM50 genes overexpressed in the luminal subtype. Similar to the clinical data, the different behaviour of Deconvolution, Guided Backpropagation, and LRP-*α*1-*β*0 calls for further studies to analyse their relevance attributions on this type of data and combining the LRP methods could lead to better results [25]. Lastly, KernelShap, Lime, Occlusion, and ShapleyValueSampling performed differently, giving more smaller attributions and varying strongly in which feature had the highest relevance attribution. This behaviour also calls for further analysis.

The differing behaviour of Deconvolution, Guided Backpropagation, and LRP-*α*1-*β*0 in the clinical and biomolecular benchmarking tasks needs to be addressed in further studies. Having inconsistent explanations in a medical setting can not only confuse clinicians and thus reduce trust in AI systems but also enhance learned biases. Additionally, there is a potential risk of diagnostic errors if the explanations misleadingly focus on irrelevant features.

As of now, BenchXAI does not yet support more complex state-of-the-art AI models, such as graph neural networks or attention-based models. This is largely due to the fact that these methods have specifically implemented XAI methods, which are not yet supported by the CaptumAI [26] platform, like GNNExplainer [61]. Including these models and their XAI methods in BenchXAI in the future will allow for even larger and more extensive benchmarking studies. Additionally, our benchmarking study currently only includes one medical image dataset, synthetic signal dataset, and biomolecular dataset. Including even more datasets of each data modality in a future study will allow for an even better comparison of XAI performance in the different bench-marking tasks. Finally, closer collaborations with domain experts in the future will allow us to further analyse and validate the explanations generated by the XAI methods to identify potential biases or even causalities.

With the ongoing advancements in the field of XAI, the future of biomedicine holds the promise of transparent, trustworthy, and personalized healthcare, enabling doctors and researchers to make informed decisions with deeper insights into complex biological systems.

## Supporting information

Full Supplementary Material

## Acknowledgements

The results shown here are in part based upon data generated by the TCGA Research Network: https://www.cancer.gov/tcga. This work is supported in part by funds from the German Ministry of Education and Research (BMBF) under grant agreement No. 01KD2208A and No. 01KD2414A (project FAIrPaCT) and by the Innovation Committee at the Federal Joint Committee NO. 01VSF20014 (project KI-Thrust). The authors gratefully acknowledge the computing time granted by the Resource Allocation Board and provided on the supercomputer Emmy/Grete at NHR-Nord@Göttingen as part of the NHR infrastructure. The calculations for this research were conducted with computing resources under the project nim00014. This work used the Scientific Compute Cluster at GWDG, the joint data center of Max Planck Society for the Advancement of Science (MPG) and University of Göttingen. In part funded by the Deutsche Forschungsgemeinschaft (DFG, German Research Foundation) – 405797229.

Figures 1 and 3 were created with BioRender.com.

## 7 Code Availablity

The BenchXAI package is available on Github: https://github.com/HauschildLab/BenchXAI.

## 8 Authors’ contributions

ACH and JMM conceptualised the project. JMM performed all programming tasks as well as analysis tasks. All authors performed review and editing.

## References

[1] Michael D. Abràmoff, Mona K. Garvin, and Milan Sonka. Retinal imaging and image analysis. IEEE Reviews in Biomedical Engineering, 3:169–208, 2010.

[2] Chirag Agarwal, Satyapriya Krishna, Eshika Saxena, Martin Pawelczyk, Nari Johnson, Isha Puri, Marinka Zitnik, and Himabindu Lakkaraju. Openxai: Towards a transparent evaluation of model explanations. Advances in neural information processing systems, 35:15784–15799, 2022.

[3] Tanvir Ahmad, Assia Munir, Sajjad Haider Bhatti, Muhammad Aftab, and Muhammad Ali Raza. Survival analysis of heart failure patients: A case study. PLOS ONE, 12(7):1–8, 07 2017.

[4] Sebastian Bach, Alexander Binder, Grégoire Montavon, Frederick Klauschen, Klaus-Robert Müller, and Wojciech Samek. On pixel-wise explanations for non-linear classifier decisions by layer-wise relevance propagation. PLOS ONE, 10(7):1–46, 07 2015.

[5] Theresa Bender, Jacqueline M Beinecke, Dagmar Krefting, Carolin Müller, Henning Dathe, Tim Seidler, Nicolai Spicher, and Anne-Christin Hauschild. Analysis of a deep learning model for 12-lead ecg classification reveals learned features similar to diagnostic criteria. IEEE Journal of Biomedical and Health Informatics, 28(4):1848–1859, 2023.

[6] Guido Bologna and Yoichi Hayashi. Qsvm: A support vector machine for rule extraction. In Advances in Computational Intelligence: 13th International Work-Conference on Artificial Neural Networks, IWANN 2015, Palma de Mallorca, Spain, June 10-12, 2015. Proceedings, Part II 13, pages 276–289. Springer, 2015.

[7] G. Bradski. The OpenCV Library. Dr. Dobb’s Journal of Software Tools, 2000.

[8] Prabir Burman. A comparative study of ordinary cross-validation, v-fold cross-validation and the repeated learning-testing methods. Biometrika, 76(3):503–514, 09 1989.

[9] Yan-Cheng Chen, Chao-Ton Su, and Taho Yang. Rule extraction from support vector machines by genetic algorithms. Neural Computing and Applications, 23:729–739, 2013.

[10] Davide Chicco and Giuseppe Jurman. The advantages of the matthews correlation coefficient (mcc) over f1 score and accuracy in binary classification evaluation. BMC genomics, 21:1–13, 2020.

[11] Davide Chicco and Giuseppe Jurman. Machine learning can predict survival of patients with heart failure from serum creatinine and ejection fraction alone. BMC Medical Informatics and Decision Making, 20, 02 2020.

[12] Wlodzislaw Duch, Rafal Adamczak, and Krzysztof Grabczewski. A new methodology of extraction, optimization and application of crisp and fuzzy logical rules. IEEE Transactions on Neural Networks, 12(2):277– 306, 2001.

[13] Jamie Duell, Xiuyi Fan, Bruce Burnett, Gert Aarts, and Shang-Ming Zhou. A comparison of explanations given by explainable artificial intelligence methods on analysing electronic health records. In 2021 IEEE EMBS International Conference on Biomedical and Health Informatics (BHI), pages 1–4. IEEE, 2021.

[14] Olive Jean Dunn. Multiple comparisons among means. Journal of the American statistical association, 56(293):52–64, 1961.

[15] Kountay Dwivedi, Ankit Rajpal, Sheetal Rajpal, Manoj Agarwal, Virendra Kumar, and Naveen Kumar. An explainable ai-driven biomarker discovery framework for non-small cell lung cancer classification. Computers in Biology and Medicine, 153:106544, 2023.

[16] Thomas Fel, Lucas Hervier, David Vigouroux, Antonin Poche, Justin Plakoo, Remi Cadene, Mathieu Chalvidal, Julien Colin, Thibaut Boissin, Louis Bethune, et al. Xplique: A deep learning explainability toolbox. *arXiv preprint arXiv:2206.04394*, 2022.

[17] Milton Friedman. The use of ranks to avoid the assumption of normality implicit in the analysis of variance. Journal of the american statistical association, 32(200):675–701, 1937.

[18] Yoichi Hayashi and Satoshi Nakano. Use of a recursive-rule extraction algorithm with j48graft to achieve highly accurate and concise rule extraction from a large breast cancer dataset. Informatics in Medicine Unlocked, 1:9–16, 2015.

[19] Anna Hedström, Philine Bommer, Kristoffer K Wickstrøm, Wojciech Samek, Sebastian Lapuschkin, and Marina M-C Höhne. The meta-evaluation problem in explainable ai: identifying reliable estimators with metaquantus. *arXiv preprint arXiv:2302.07265*, 2023.

[20] Chang Hu, Chao Gao, Tianlong Li, Chang Liu, and Zhiyong Peng. Explainable artificial intelligence model for mortality risk prediction in the intensive care unit: a derivation and validation study. Postgraduate Medical Journal, 100(1182):219–227, 01 2024.

[21] Shujun Huang, Leigh Murphy, and Wayne Xu. Genes and functions from breast cancer signatures. BMC cancer, 18:1–15, 2018.

[22] Md Abdul Kadir, Amir Mosavi, and Daniel Sonntag. Evaluation metrics for xai: A review, taxonomy, and practical applications. In 2023 IEEE 27th International Conference on Intelligent Engineering Systems (INES), pages 000111–000124. IEEE, 2023.

[23] Md. Sarwar Kamal, Aden Northcote, Linkon Chowdhury, Nilanjan Dey, Rubén González Crespo, and Enrique Herrera-Viedma. Alzheimer’s patient analysis using image and gene expression data and explainable-ai to present associated genes. IEEE Transactions on Instrumentation and Measurement, 70:1–7, 2021.

[24] Eamonn Keogh and Abdullah Mueen. *Curse of Dimensionality*, pages 314–315. Springer US, Boston, MA, 2017.

[25] Maximilian Kohlbrenner, Alexander Bauer, Shinichi Nakajima, Alexander Binder, Wojciech Samek, and Sebastian Lapuschkin. Towards best practice in explaining neural network decisions with lrp. 2020 *International Joint Conference on Neural Networks (IJCNN)*, pages 1–7, 2019.

[26] Narine Kokhlikyan, Vivek Miglani, Miguel Martin, Edward Wang, Bilal Alsallakh, Jonathan Reynolds, Alexander Melnikov, Natalia Kliushkina, Carlos Araya, Siqi Yan, and Orion Reblitz-Richardson. Captum: A unified and generic model interpretability library for pytorch, 2020.

[27] Johann Laux, Sandra Wachter, and Brent Mittelstadt. Trustworthy artificial intelligence and the european union ai act: On the conflation of trustworthiness and acceptability of risk. Regulation & Governance, 18(1):3–32, 2024.

[28] Luca Longo, Mario Brcic, Federico Cabitza, Jaesik Choi, Roberto Confalonieri, Javier Del Ser, Riccardo Guidotti, Yoichi Hayashi, Francisco Herrera, Andreas Holzinger, et al. Explainable artificial intelligence (xai) 2.0: A manifesto of open challenges and interdisciplinary research directions. Information Fusion, 106:102301, 2024.

[29] Samual MacDonald, Kaiah Steven, and Maciej Trzaskowski. Interpretable ai in healthcare: Enhancing fairness, safety, and trust. In *Artificial Intelligence in Medicine: Applications*, Limitations and Future Directions, pages 241–258. Springer, 2022.

[30] Olvi L Mangasarian and William H Wolberg. Cancer diagnosis via linear programming. Technical report, University of Wisconsin-Madison Department of Computer Sciences, 1990.

[31] Jacqueline Michelle Metsch, Anna Saranti, Alessa Angerschmid, Bastian Pfeifer, Vanessa Klemt, Andreas Holzinger, and Anne-Christin Hauschild. Clarus: An interactive explainable ai platform for manual counterfactuals in graph neural networks. Journal of Biomedical Informatics, 150:104600, 2024.

[32] Grégoire Montavon, Alexander Binder, Sebastian Lapuschkin, Wojciech Samek, and Klaus-Robert Müller. Layer-Wise Relevance Propagation: An Overview, pages 193–209. Springer International Publishing, Cham, 2019.

[33] Grégoire Montavon, Sebastian Lapuschkin, Alexander Binder, Wojciech Samek, and Klaus-Robert Müller. Explaining nonlinear classification decisions with deep taylor decomposition. Pattern Recognition, 65:211–222, May 2017.

[34] Detlef Nauck and Rudolf Kruse. Obtaining interpretable fuzzy classification rules from medical data. Artificial intelligence in medicine, 16(2):149–169, 1999.

[35] Youngjun Park, Nils P Muttray, and Anne-Christin Hauschild. Species-agnostic transfer learning for cross-species transcriptomics data integration without gene orthology. Briefings in Bioinformatics, 25(2):bbae004, 02 2024.

[36] Joel S Parker, Michael Mullins, Maggie CU Cheang, Samuel Leung, David Voduc, Tammi Vickery, Sherri Davies, Christiane Fauron, Xiaping He, Zhiyuan Hu, et al. Supervised risk predictor of breast cancer based on intrinsic subtypes. Journal of clinical oncology, 27(8):1160–1167, 2009.

[37] Adam Paszke, Sam Gross, Francisco Massa, Adam Lerer, James Bradbury, Gregory Chanan, Trevor Killeen, Zeming Lin, Natalia Gimelshein, Luca Antiga, Alban Desmaison, Andreas Kopf, Edward Yang, Zachary DeVito, Martin Raison, Alykhan Tejani, Sasank Chilamkurthy, Benoit Steiner, Lu Fang, Junjie Bai, and Soumith Chintala. Pytorch: An imperative style, high-performance deep learning library. In Advances in Neural Information Processing Systems 32, pages 8024–8035. Curran Associates, Inc., 2019.

[38] F. Pedregosa, G. Varoquaux, A. Gramfort, V. Michel, B. Thirion, O. Grisel, M. Blondel, P. Prettenhofer, R. Weiss, V. Dubourg, J. Vanderplas, A. Passos, D. Cournapeau, M. Brucher, M. Perrot, and E. Duchesnay. Scikit-learn: Machine learning in Python. Journal of Machine Learning Research, 12:2825–2830, 2011.

[39] Prasanna Porwal, Samiksha Pachade, Ravi Kamble, Manesh Kokare, Girish Deshmukh, Vivek Sahasrabuddhe, and Fabrice Meriaudeau. Indian diabetic retinopathy image dataset (idrid): a database for diabetic retinopathy screening research. Data, 3(3):25, 2018.

[40] Prasanna Porwal, Samiksha Pachade, Manesh Kokare, Girish Deshmukh, Jaemin Son, Woong Bae, Lihong Liu, Jianzong Wang, Xinhui Liu, Liangxin Gao, TianBo Wu, Jing Xiao, Fengyan Wang, Baocai Yin, Yunzhi Wang, Gopichandh Danala, Linsheng He, Yoon Ho Choi, Yeong Chan Lee, Sang-Hyuk Jung, Zhongyu Li, Xiaodan Sui, Junyan Wu, Xiaolong Li, Ting Zhou, Janos Toth, Agnes Baran, Avinash Kori, Sai Saketh Chennamsetty, Mohammed Safwan, Varghese Alex, Xingzheng Lyu, Li Cheng, Qinhao Chu, Pengcheng Li, Xin Ji, Sanyuan Zhang, Yaxin Shen, Ling Dai, Oindrila Saha, Rachana Sathish, Tânia Melo, Teresa Araújo, Balazs Harangi, Bin Sheng, Ruogu Fang, Debdoot Sheet, Andras Hajdu, Yuanjie Zheng, Ana Maria Mendonça, Shaoting Zhang, Aurélio Campilho, Bin Zheng, Dinggang Shen, Luca Giancardo, Gwenolé Quellec, and Fabrice Mériaudeau. Idrid: Diabetic retinopathy – segmentation and grading challenge. Medical Image Analysis, 59:101561, 2020.

[41] Elias Reichel and David Salz. Diabetic retinopathy screening. Managing Diabetic Eye Disease in Clinical Practice, pages 25–38, 2015.

[42] Marco Ribeiro, Sameer Singh, and Carlos Guestrin. “why should I trust you?”: Explaining the predictions of any classifier. In Proceedings of the 2016 Conference of the North American Chapter of the Association for Computational Linguistics: Demonstrations, pages 97–101, San Diego, California, June 2016. Association for Computational Linguistics.

[43] Wojciech Samek, Alexander Binder, Grégoire Montavon, Sebastian Lapuschkin, and Klaus-Robert Müller. Evaluating the visualization of what a deep neural network has learned. IEEE Transactions on Neural Networks and Learning Systems, 28(11):2660–2673, 2017.

[44] M Scott, Lee Su-In, et al. A unified approach to interpreting model predictions. Advances in neural information processing systems, 30:4765–4774, 2017.

[45] L. S. Shapley and Martin Shubik. A method for evaluating the distribution of power in a committee system. American Political Science Review, 48(3):787–792, 1954.

[46] Lloyd S. Shapley. A Value for N-Person Games. RAND Corporation, Santa Monica, CA, 1952.

[47] Sumeet Shinde, Tanay Chougule, Jitender Saini, and Madhura Ingalhalikar. Hr-cam: Precise localization of pathology using multi-level learning in cnns. In Dinggang Shen, Tianming Liu, Terry M. Peters, Lawrence H. Staib, Caroline Essert, Sean Zhou, Pew-Thian Yap, and Ali Khan, editors, *M*edical Image Computing and Computer Assisted Intervention – MICCAI 2019, pages 298–306, Cham, 2019. Springer International Publishing.

[48] Avanti Shrikumar, Peyton Greenside, and Anshul Kundaje. Learning important features through propagating activation differences. In Proceedings of the 34th International Conference on Machine Learning - Volume 70, ICML’17, page 3145–3153. JMLR.org, 2017.

[49] Avanti Shrikumar, Peyton Greenside, Anna Shcherbina, and Anshul Kundaje. Not just a black box: Learning important features through propagating activation differences, 2017.

[50] Siegel Sidney. Nonparametric statistics for the behavioral sciences. The Journal of Nervous and Mental Disease, 125(3):497, 1957.

[51] Karen Simonyan, Andrea Vedaldi, and Andrew Zisserman. Deep inside convolutional networks: Visualising image classification models and saliency maps, 2014.

[52] J.T. Springenberg, A. Dosovitskiy, T. Brox, and M. Riedmiller. Striving for simplicity: The all convolutional net. In ICLR (workshop track), 2015.

[53] P Naga Srinivasu, G Jaya Lakshmi, Abhishek Gudipalli, Sujatha Canavoy Narahari, Jana Shafi, Marcin Woźniak, and Muhammad Fazal Ijaz. Xai-driven catboost multi-layer perceptron neural network for analyzing breast cancer. Scientific Reports, 14(1):28674, 2024.

[54] Erik Strumbelj and Igor Kononenko. An efficient explanation of individual classifications using game theory. J. Mach. Learn. Res., 11:1–18, 2010.

[55] Mukund Sundararajan, Ankur Taly, and Qiqi Yan. Axiomatic attribution for deep networks, 2017.

[56] Hugues Turbé, Mina Bjelogrlic, Christian Lovis, and Gianmarco Mengaldo. Evaluation of post-hoc interpretability methods in time-series classification. Nature Machine Intelligence, 5(3):250–260, 2023.

[57] Guido Van Rossum and Fred L Drake Jr. *Python reference manual*. Centrum voor Wiskunde en Informatica Amsterdam, 1995.

[58] WIlliam Wolberg. Breast Cancer Wisconsin (Original). UCI Machine Learning Repository, 1992. DOI: 10.24432/C5HP4Z.

[59] Lihteh Wu, Priscilla Fernandez-Loaiza, Johanna Sauma, Erick Hernandez-Bogantes, and Marissé Masis. Classification of diabetic retinopathy and diabetic macular edema. World journal of diabetes, 4(6):290, 2013.

[60] Chih-Kuan Yeh, Cheng-Yu Hsieh, Arun Suggala, David I Inouye, and Pradeep K Ravikumar. On the (in) fidelity and sensitivity of explanations. Advances in neural information processing systems, 32, 2019.

[61] Rex Ying, Dylan Bourgeois, Jiaxuan You, Marinka Zitnik, and Jure Leskovec. Gnnexplainer: Generating explanations for graph neural networks. Advances in neural information processing systems, 32:9240, 2019.

[62] Matthew D. Zeiler and Rob Fergus. Visualizing and understanding convolutional networks. In David Fleet, Tomas Pajdla, Bernt Schiele, and Tinne Tuytelaars, editors, Computer Vision – ECCV 2014, pages 818–833, Cham, 2014. Springer International Publishing.

[63] Luisa M Zintgraf, Taco S Cohen, Tameem Adel, and Max Welling. Visualizing deep neural network decisions: Prediction difference analysis. In International Conference on Learning Representations, 2017.

