## Supplementary material for "BenchXAI: Comprehensive Benchmarking of Post-hoc Explainable AI Methods on Multi-Modal Biomedical Data": Full Supplementary Material

### 1 Supplementary

#### 1.1 Data

| Feature | Description |
|---|---|
| CT | Clump thickness |
| UCSI | Uniformity of cell size |
| UCSH | Uniformity of cell shape |
| MA | Marginal adhesion |
| SECS | Single epithelial cell size |
| BN | Bare nuclei |
| BC | Bland chromatin |
| NN | Normal nucleoli |
| M | Mitoses |
| Class | Benign or malignant tumor |

Table S1: List of all features in the Wisconsin Breast Cancer Dataset (WBCD) dataset.

| Feature | Description |
|---|---|
| Age | Age of the patient |
| A | Aneamia: Decrease of red blood cells or hemoglobin |
| HBP | High blood pressure: If a patient has hypertension |
| CPK | Creatinine Phosphokinase: Level of the CPK enzyme in the blood |
| D | Diabetes: If the patient has diabetes |
| EF | Ejection fraction: Percentage of blood leaving the heart per contraction |
| Sex | Sex of a patient |
| P | Platelets in the blood |
| SC | Level of creatinine in the blood |
| SS | Level of sodium in the blood |
| S | If the patient smokes |
| Time | Length of follow-up period |
| Target | Death Event: If the patient died during the follow-up period |

Table S2: List of all features for the Heart Disease Dataset (Heart).

#### 1.2 Results

##### 1.2.1 Task 1: XAI Benchmark for Medical Data

| | | NN | | | LogReg | | | RF | | |
|---|---|---|---|---|---|---|---|---|---|---|
| | | Train | Val | Test | Train | Val | Test | Train | Val | Test |
| | AUROC | 1.0 | 1.0 | 1.0 | 1.0 | 1.0 | 1.0 | 1.0 | 1.0 | 1.0 |
| WBCD | AUPRC | 1.0 | 1.0 | 1.0 | 1.0 | 1.0 | 1.0 | 1.0 | 1.0 | 1.0 |
| | MCC | 0.838 | 0.82 | 0.718 | 0.994 | 0.975 | 0.942 | 1.0 | 1.0 | 0.961 |
| | AUROC | 0.951 | 0.738 | 0.743 | 0.906 | 0.871 | 0.809 | 1.0 | 0.888 | 0.911 |
| Heart | AUPRC | 0.89 | 0.605 | 0.771 | 0.767 | 0.697 | 0.788 | 1.0 | 0.732 | 0.904 |
| | MCC | 0.708 | 0.339 | 0.366 | 0.508 | 0.419 | 0.383 | 1.0 | 0.609 | 0.566 |
| | AUROC | 1.0 | 1.0 | 1.0 | 1.0 | 1.0 | 1.0 | 1.0 | 1.0 | 1.0 |
| BRCA | AUPRC | 1.0 | 1.0 | 1.0 | 1.0 | 1.0 | 1.0 | 1.0 | 1.0 | 1.0 |
| | MCC | 0.838 | 0.82 | 0.718 | 0.994 | 0.975 | 0.942 | 1.0 | 1.0 | 0.961 |

Table S3: Performance of neural network (NN), logistic regression (LogReg), and random forest (RF) on training, validation, and test set on the WBCD, Heart, and BRCA datasets. Performance is shown as the median over all 100 iteration according to the area under the receiver operator curve (AUROC), the area under the precision recall curve (AUPRC) and the matthews correlation coefficient (MCC).

| | | NN | | |
|---|---|---|---|---|
| | | Train | Val | Test |
| Signal | AUROC | 1.0 | 0.999 | 0.999 |
| | AUPRC | 1.0 | 0.999 | 0.999 |
| | MCC | 0.954 | 0.938 | 0.932 |
| IDRiD | AUROC | 1.0 | 0.989 | 0.997 |
| | AUPRC | 1.0 | 0.996 | 0.997 |
| | MCC | 1.0 | 0.887 | 0.905 |

Table S4: Performance of neural network (NN) on training, validation, and test set of Signal data and IDRiD data. Performance is shown as the median over all 100 iteration according to the area under the receiver operator curve (AUROC), the area under the precision recall curve (AUPRC) and the matthews correlation coefficient (MCC).

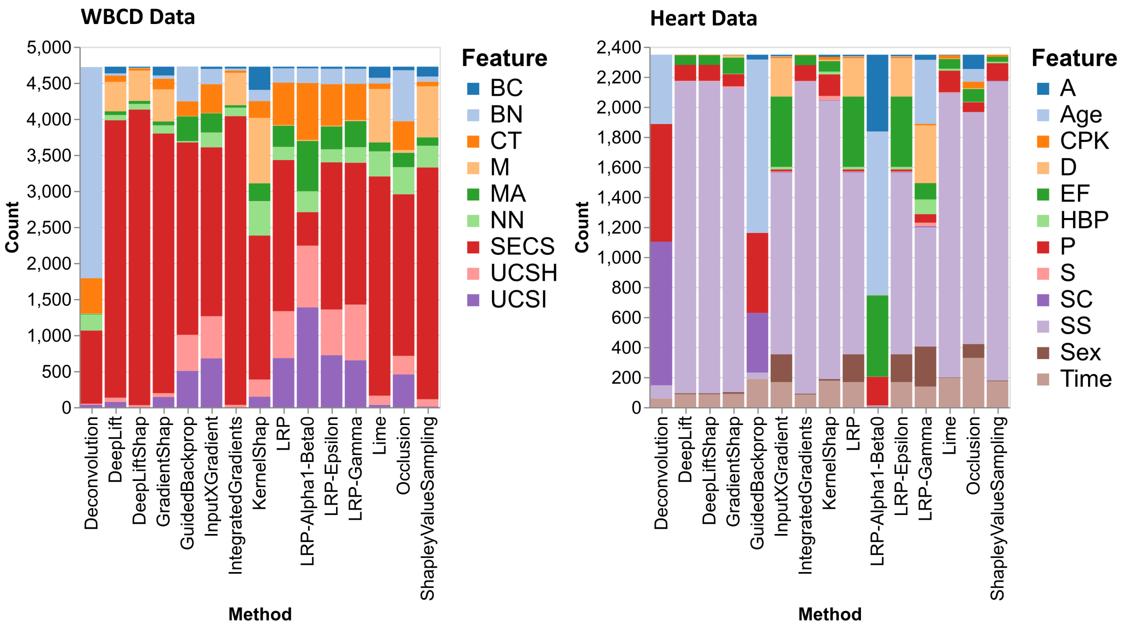


Figure S1: Stacked bar plots for the WBCD dataset (left) and the Heart dataset (right) that show how often each feature had the lowest negative attribution for all XAI methods and for all correct classified class 1 test samples over all 100 iterations.

| | Age | A | CPK | D | EF | HBP | P | SC | SS | Sex | S | Time | ∑ |
|---|---|---|---|---|---|---|---|---|---|---|---|---|---|
| Deconvolution | 0.0 | 0.0 | 0.0 | 0.0 | 0.0 | 0.0 | 0.0 | 0.0 | 0.0 | 0.0 | 0.0 | 0.0 | 0 |
| DeepLift | 0.0 | **1.0** | **1.0** | **1.0** | 0.0 | **1.0** | 0.0 | **1.0** | 0.0 | 0.0 | **1.0** | **0.09** | 7 |
| DeepLiftShap | 0.0 | **1.0** | **1.0** | **1.0** | 0.0 | **1.0** | 0.0 | **1.0** | 0.0 | 0.0 | **1.0** | **0.09** | 7 |
| GradientShap | 0.0 | **1.0** | **1.0** | **1.0** | 0.0 | **1.0** | 0.0 | 0.01 | 0.0 | 0.0 | **1.0** | **1.0** | 6 |
| GuidedBackprop | 0.0 | 0.0 | **1.0** | 0.0 | 0.0 | **1.0** | 0.0 | **1.0** | 0.0 | 0.0 | 0.0 | 0.0 | 3 |
| InputXGradient | **1.0** | **1.0** | 0.0 | **1.0** | **1.0** | **1.0** | 0.0 | **1.0** | **1.0** | 0.0 | 0.0 | **1.0** | 8 |
| IntegratedGradients | 0.0 | **1.0** | **1.0** | **1.0** | 0.0 | **1.0** | 0.0 | **1.0** | 0.0 | 0.0 | **1.0** | 0.04 | 6 |
| KernelShap | 0.0 | **1.0** | **1.0** | **1.0** | 0.0 | **0.89** | **0.38** | **1.0** | 0.0 | 0.0 | **1.0** | **1.0** | 8 |
| LRP | **1.0** | **1.0** | 0.0 | **1.0** | **1.0** | **1.0** | 0.0 | **1.0** | **1.0** | 0.0 | 0.0 | **1.0** | 8 |
| LRP-Alpha1-Beta0 | 0.0 | **1.0** | 0.01 | **1.0** | 0.0 | **1.0** | 0.0 | **0.22** | 0.0 | 0.0001 | **1.0** | 0.0002 | 5 |
| LRP-Epsilon | **1.0** | **1.0** | 0.0 | **1.0** | **1.0** | **1.0** | 0.0 | **1.0** | **1.0** | 0.0 | 0.0 | **1.0** | 8 |
| LRP-Gamma | 0.0 | **1.0** | **1.0** | **1.0** | 0.0 | **1.0** | **1.0** | **1.0** | **1.0** | **1.0** | **1.0** | **1.0** | 10 |
| Lime | 0.0 | **1.0** | **1.0** | **1.0** | 0.0 | 0.0001 | 0.0 | **1.0** | 0.0 | 0.0 | **1.0** | **1.0** | 6 |
| Occlusion | **0.12** | **1.0** | 0.0 | **1.0** | **1.0** | **1.0** | 0.0 | **1.0** | **1.0** | **1.0** | **1.0** | **1.0** | 10 |
| ShapleyValueSampling | 0.0 | **1.0** | 0.04 | **1.0** | 0.0 | 0.0 | 0.0 | **1.0** | 0.0 | 0.0 | **1.0** | **1.0** | 5 |
| ∑ | 4 | 13 | 8 | 13 | 4 | 12 | 2 | 13 | 5 | 2 | 10 | 11 | 97 |

Table S5: Heart data: Bonferroni corrected p-values of repeated Friedman tests for each feature and XAI method. Test were performed on all 10 class 1 samples that were correctly classified over all 100 iterations. All NOT significant p-values (>0.05) are highlighted in green. These indicate that $H_0$ : „No significant differences in ranked attributions of XAI Method between all 100 iterations." is not rejected.

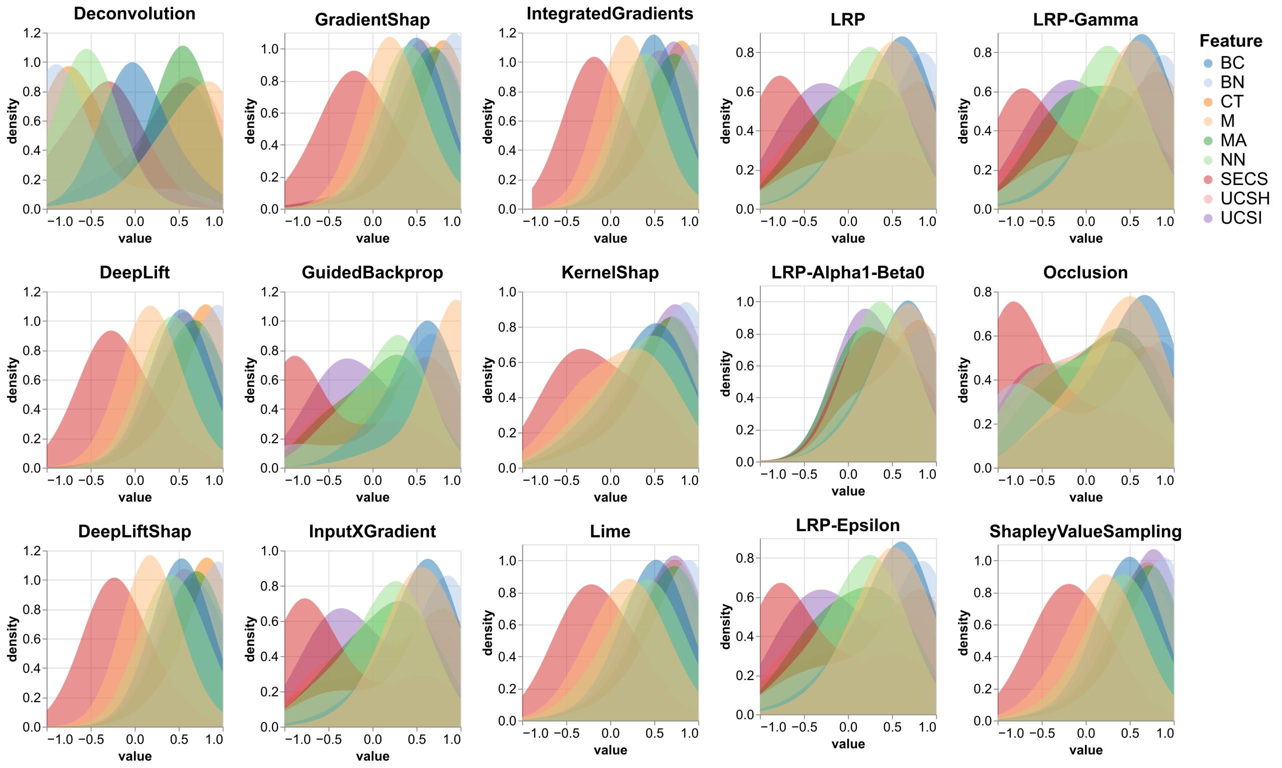


Figure S2: Stacked density plots for the WBCD dataset that show the distribution of relevance attributions for each feature and each XAI methods for all correct classified class 1 test samples over all 100 iterations.

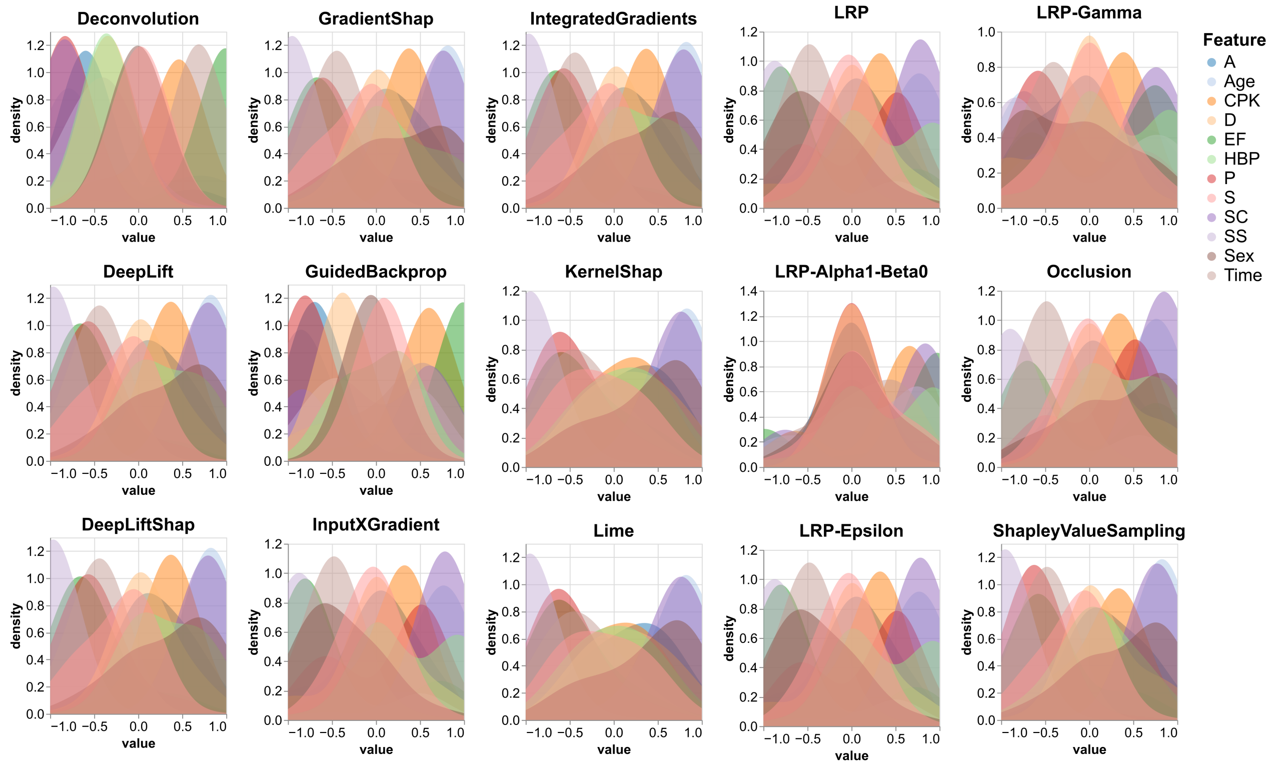


Figure S3: Stacked density plots for the Heart dataset that show the distribution of relevance attributions for each feature and each XAI methods for all correct classified class 1 test samples over all 100 iterations.

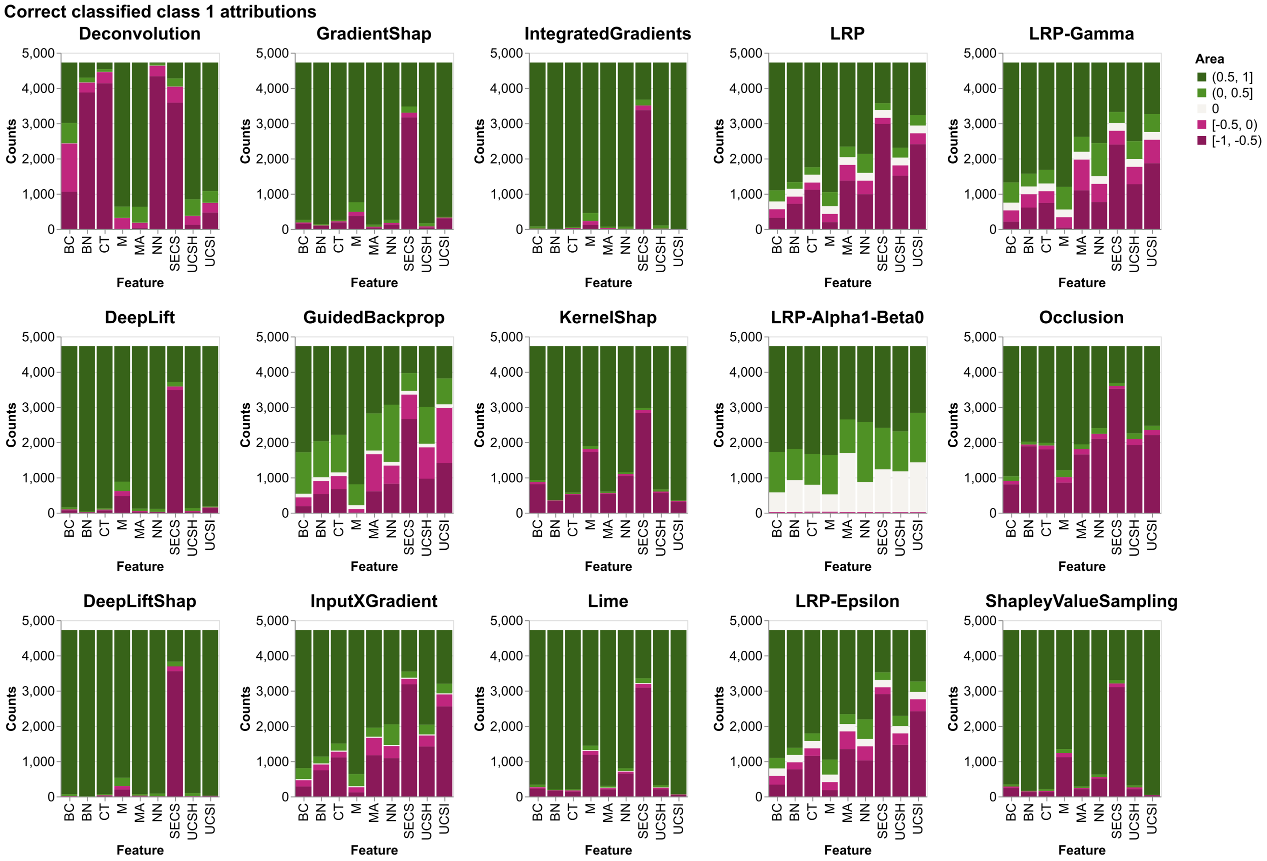


Figure S4: Stacked bar plots for the WBCD dataset that show counts of negative, positive and zero attributions for all correct classified class 1 samples and for all XAI methods.

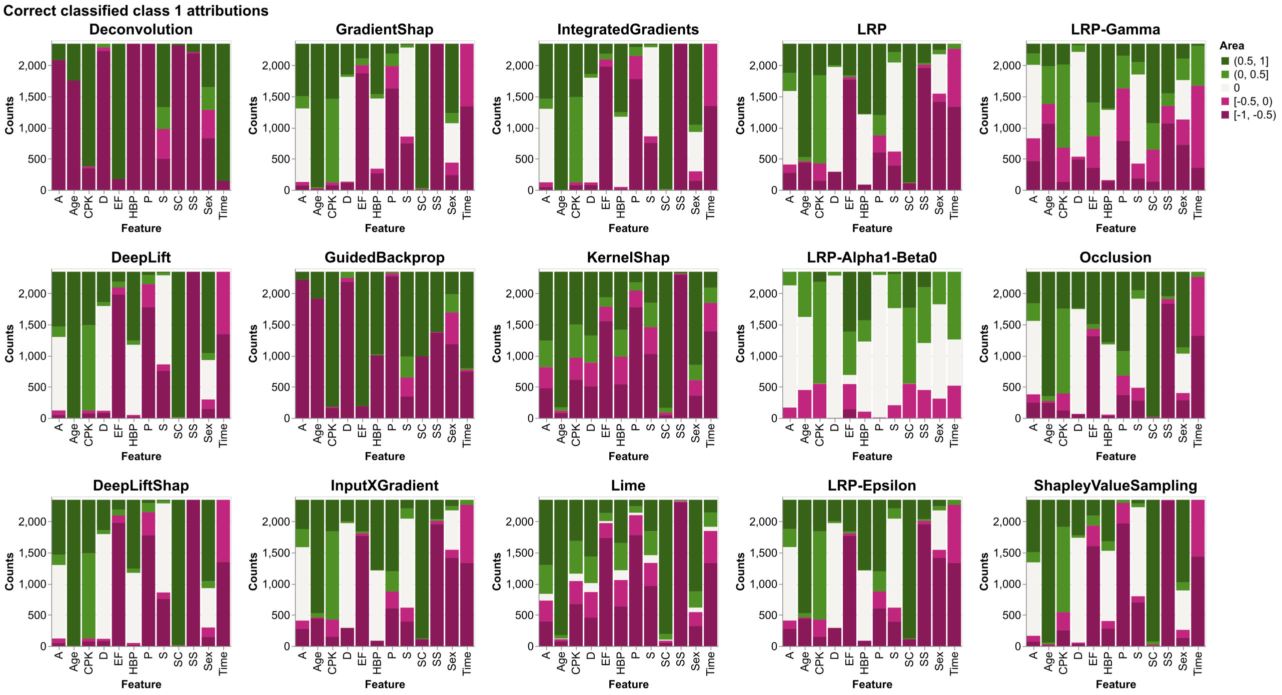


Figure S5: Stacked bar plots for the Heart dataset that show counts of negative, positive and zero attributions for all correct classified class 1 samples and for all XAI methods.

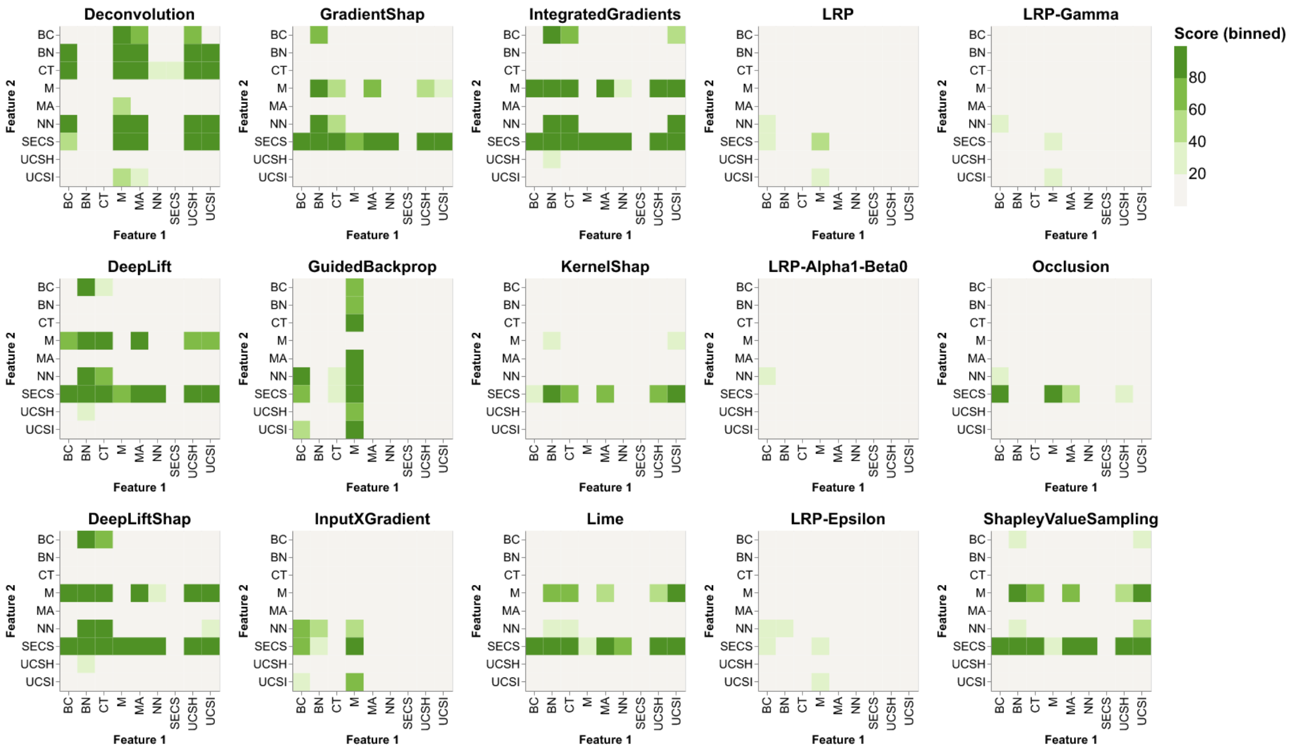


Figure S6: Sum of significant p-values (<0.05) when performing a one sided Wilcoxon test between all features for all 100 iterations for all XAI methods for the WBCD dataset. A significant p-value indicates the rejection of $H_0$ : median(Feature 1 - Feature 2) = 0 in favor of $H_1$ : median(Feature 1 - Feature 2) > 0, indicating that feature 1 is significantly larger than feature 2. P-values were corrected using Bonferroni correction to account for multiple testing.

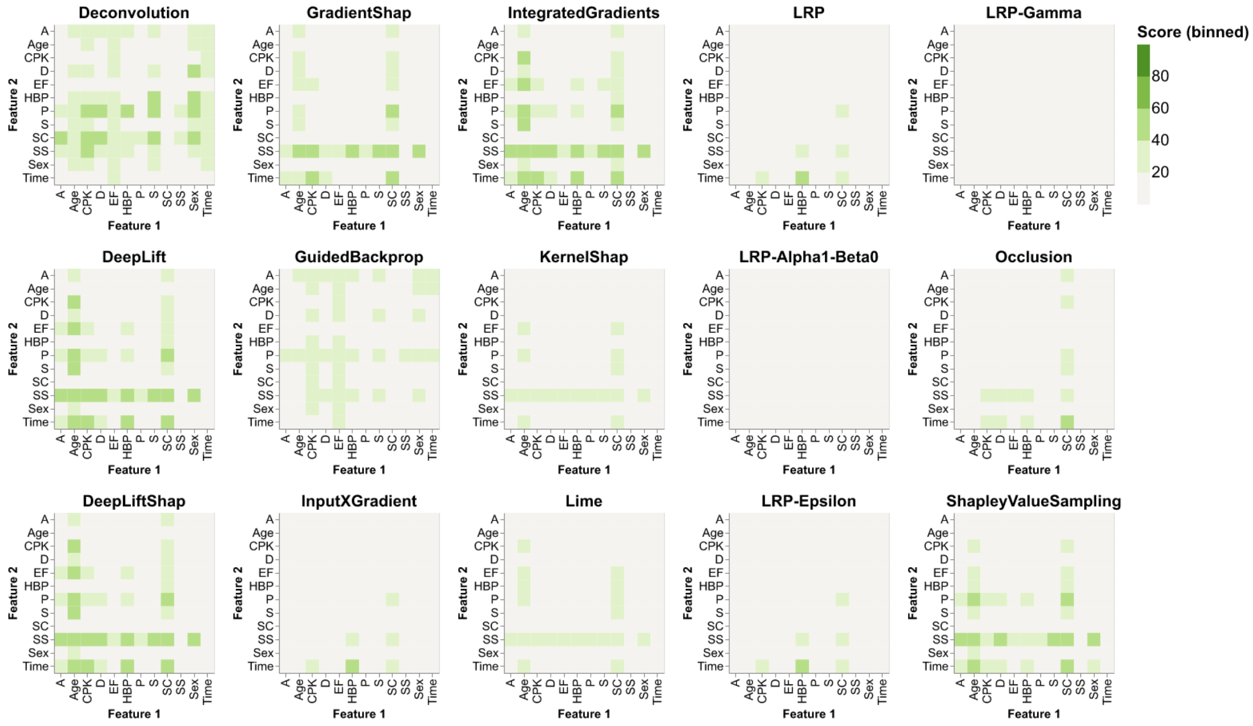


Figure S7: Sum of significant p-values (<0.05) when performing a one sided Wilcoxon test between all features for all 100 iterations for all XAI methods for the Heart dataset. A significant p-value indicates the rejection of $H_0$ : median(Feature 1 - Feature 2) = 0 in favor of $H_1$ : median(Feature 1 - Feature 2) > 0, indicating that feature 1 is significantly larger than feature 2. P-values were corrected using Bonferroni correction to account for multiple testing.

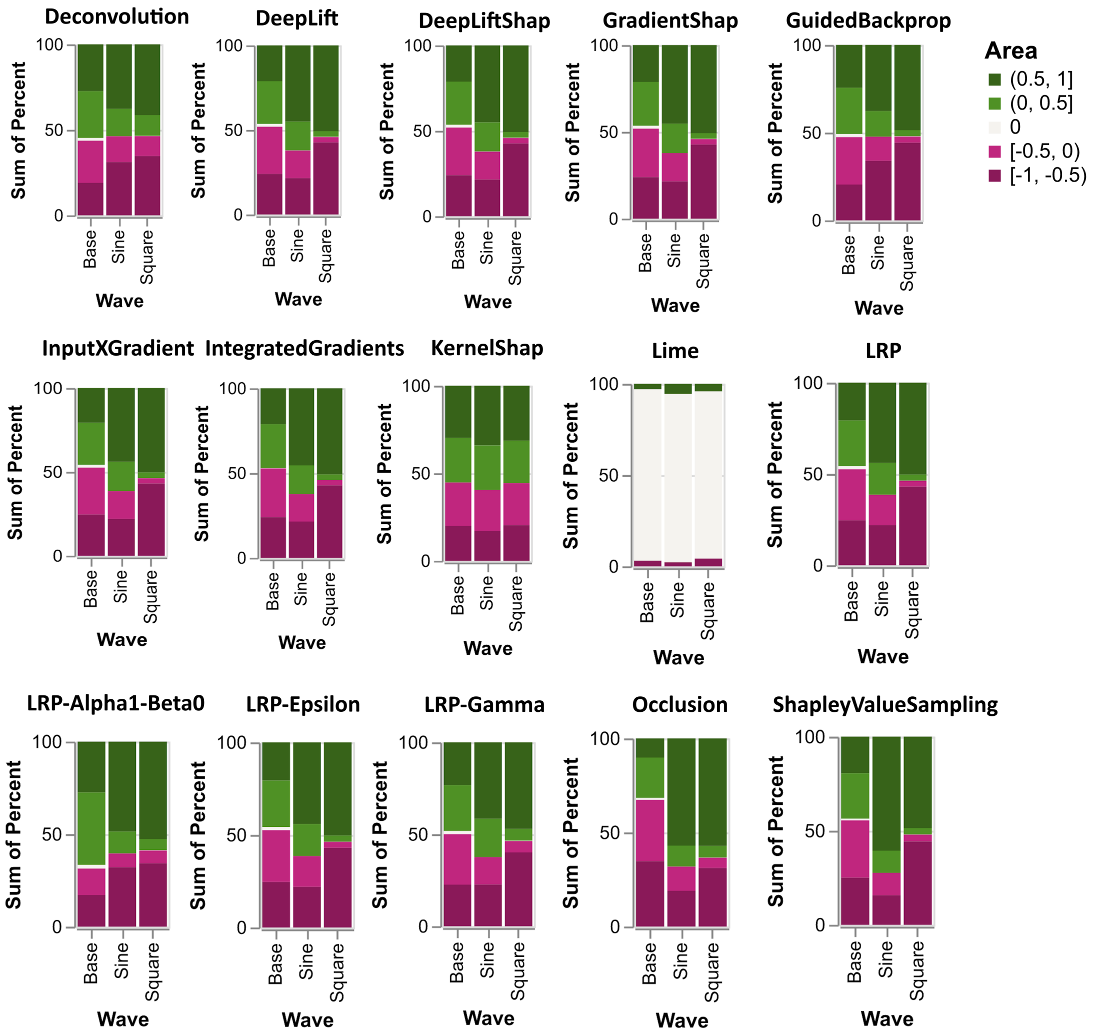


Figure S8: Stacked bar plots for the Signal dataset that show counts of negative, positive and zero attributions for all correct classified class 1 samples and for all XAI methods.

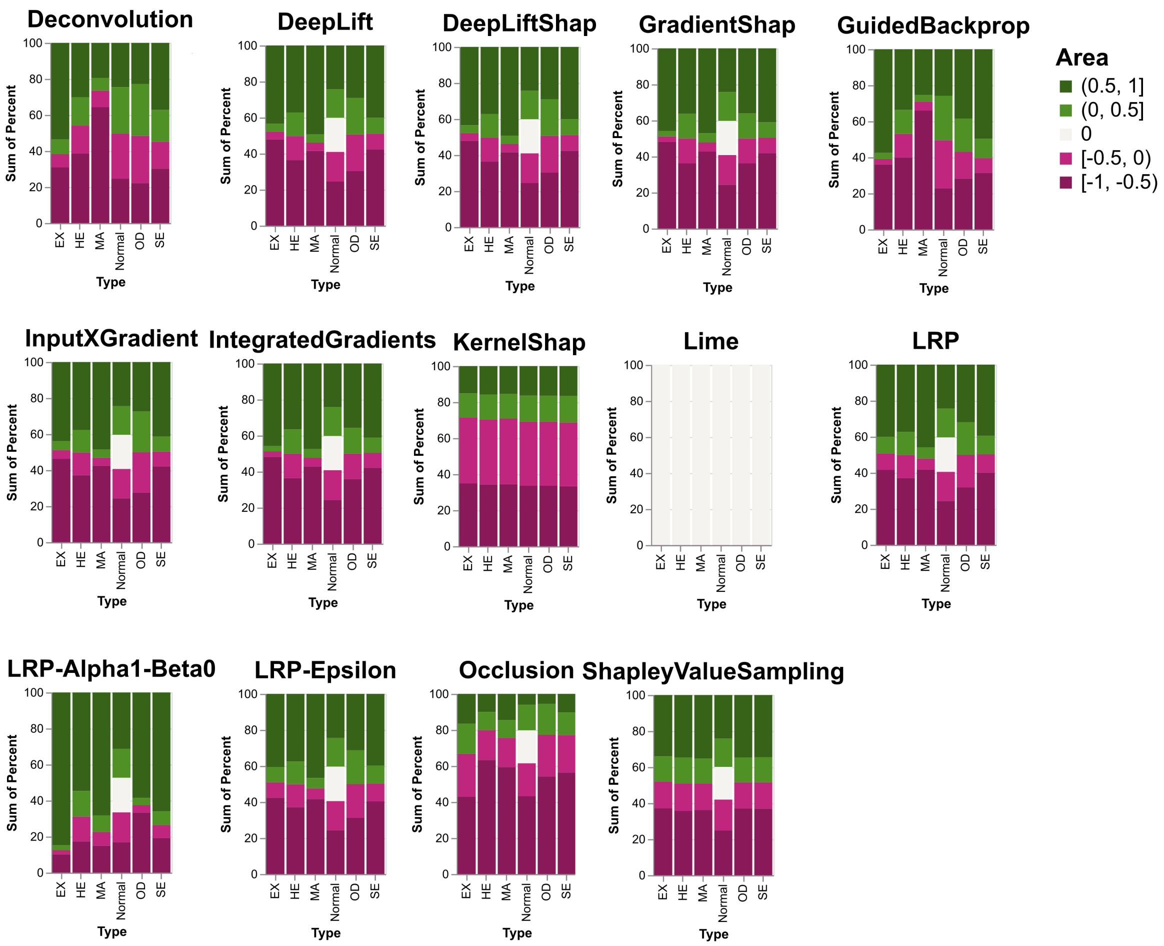


Figure S9: Stacked bar plots for the IDRiD dataset that show counts of negative, positive and zero attributions for all correct classified class 1 samples and for all XAI methods.

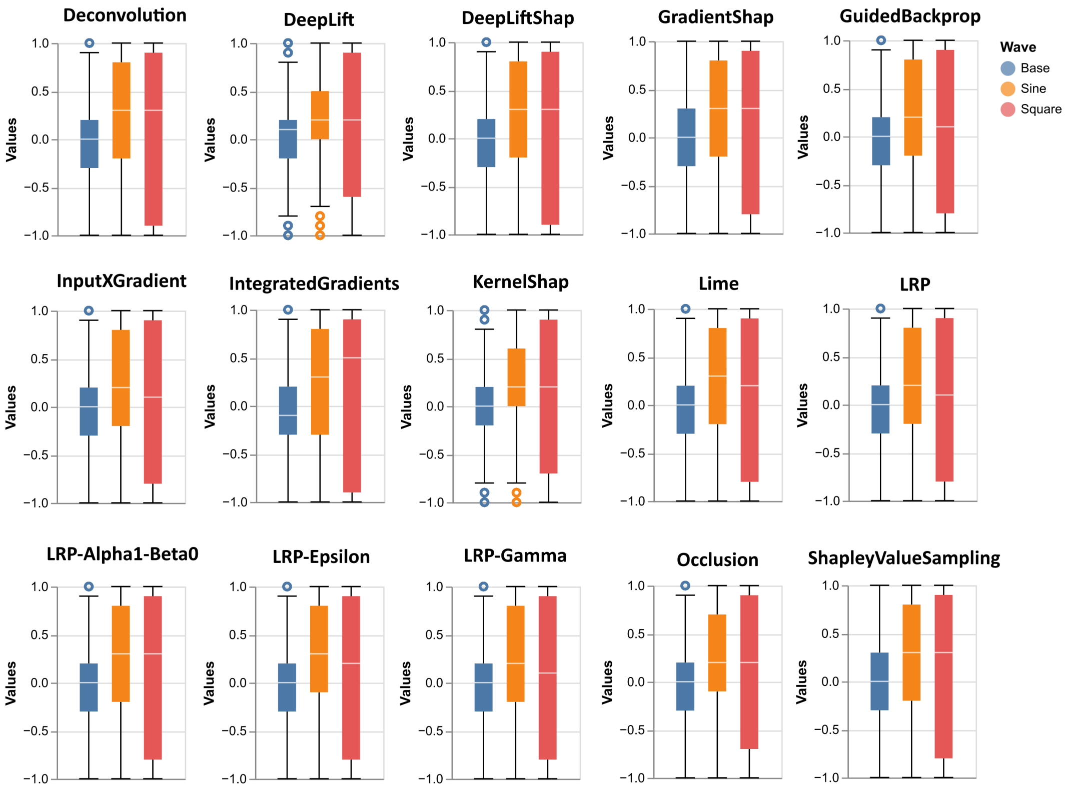


Figure S10: Boxplots of relevance attributions for all correct classified class 1 test samples of the Signal dataset for all XAI methods.

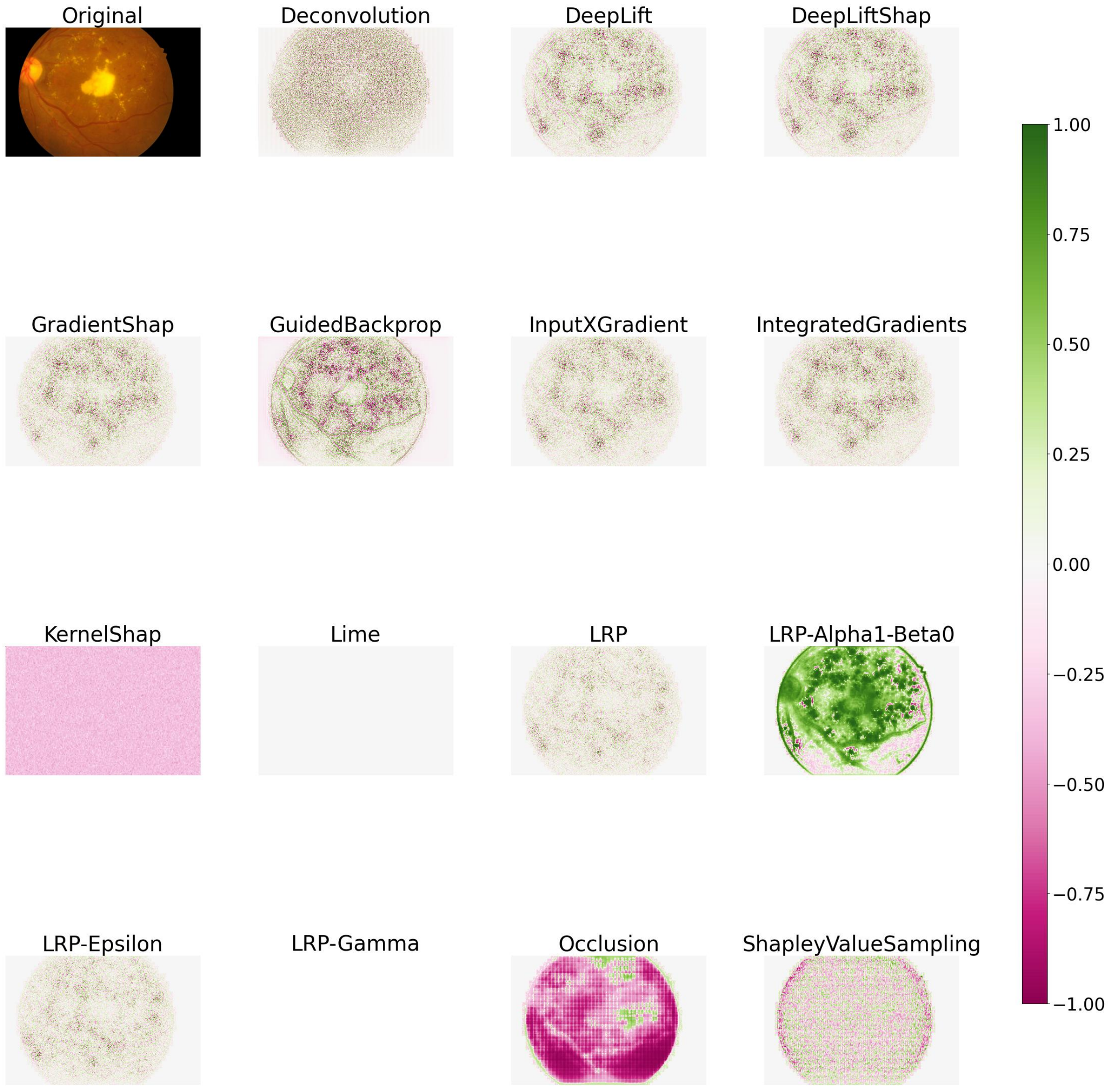


Figure S11: Original image and median relevance attributions for all XAI Methods. Medians are taken over all 100 iterations.

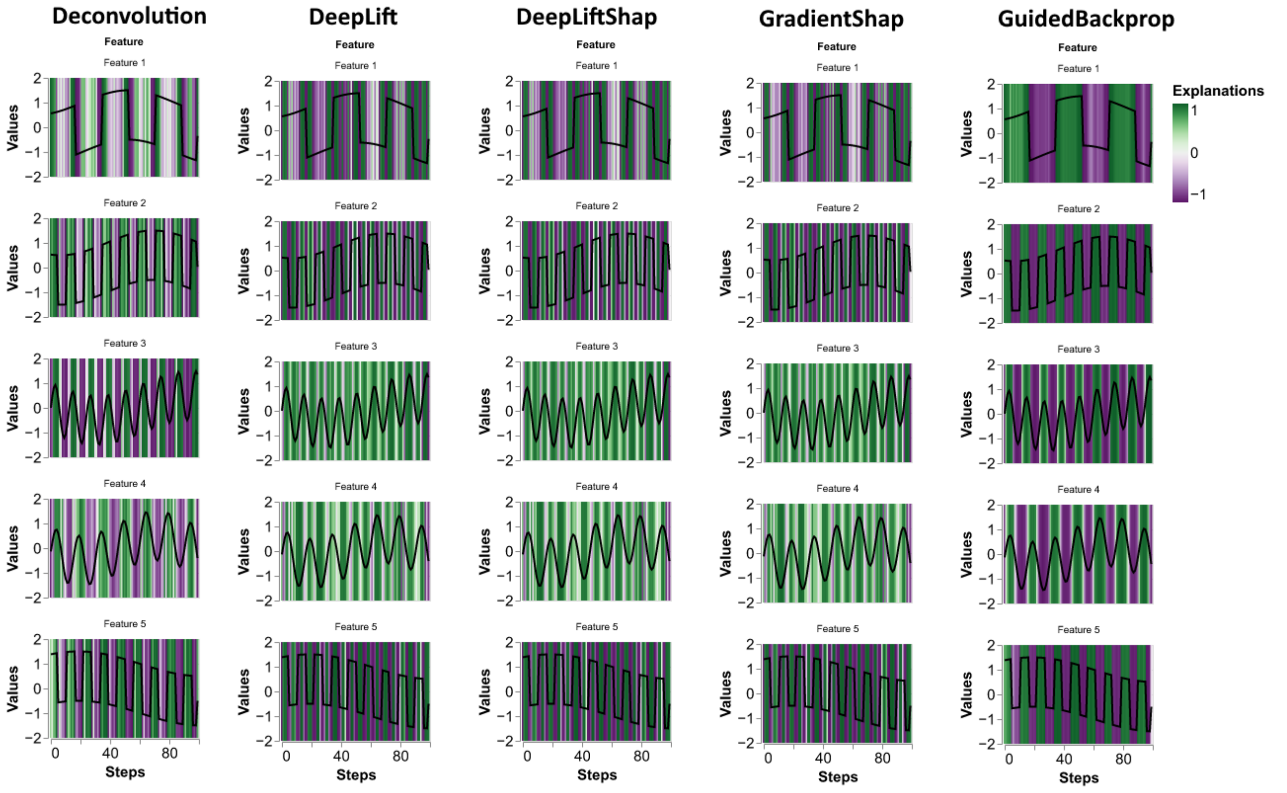


Figure S12: Median relevance attributions over all 100 iterations for one random sample for all XAI methods (Part 1).

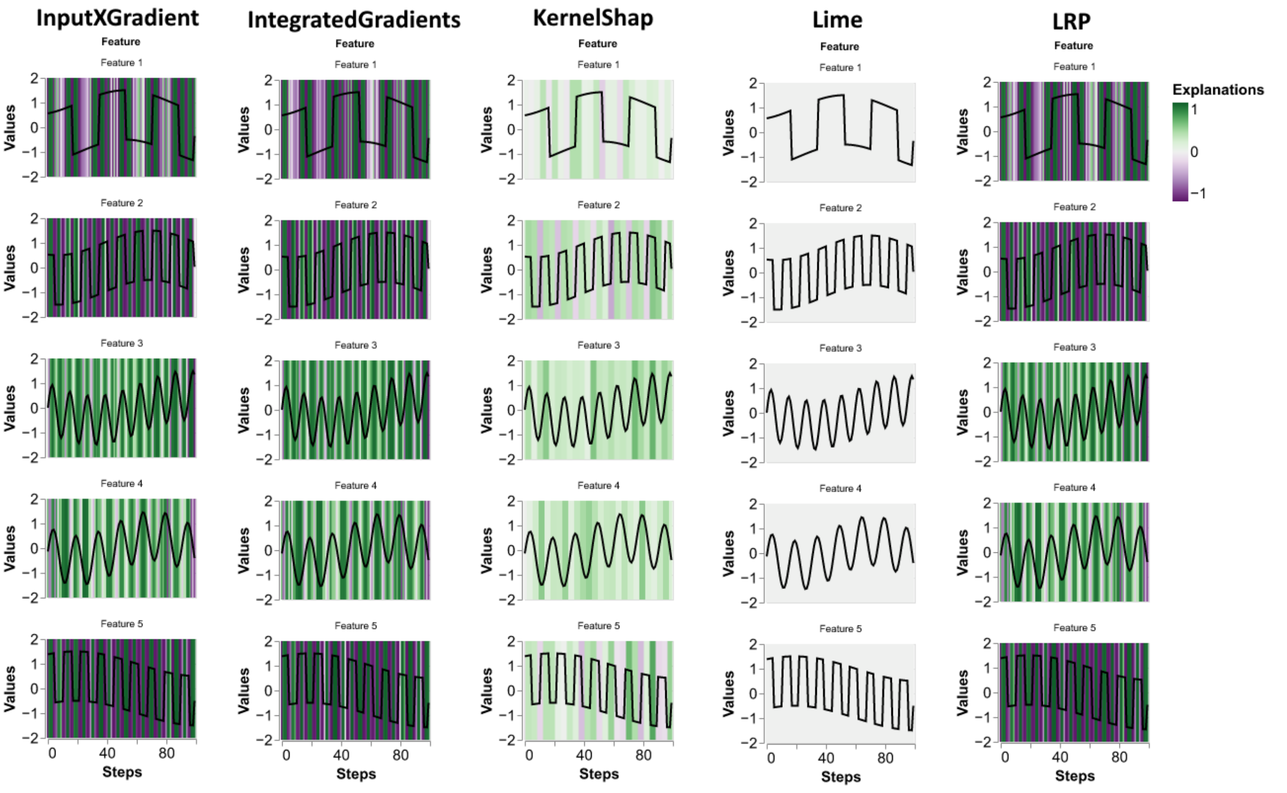


Figure S13: Median relevance attributions over all 100 iterations for one random sample for all XAI methods (Part 2).

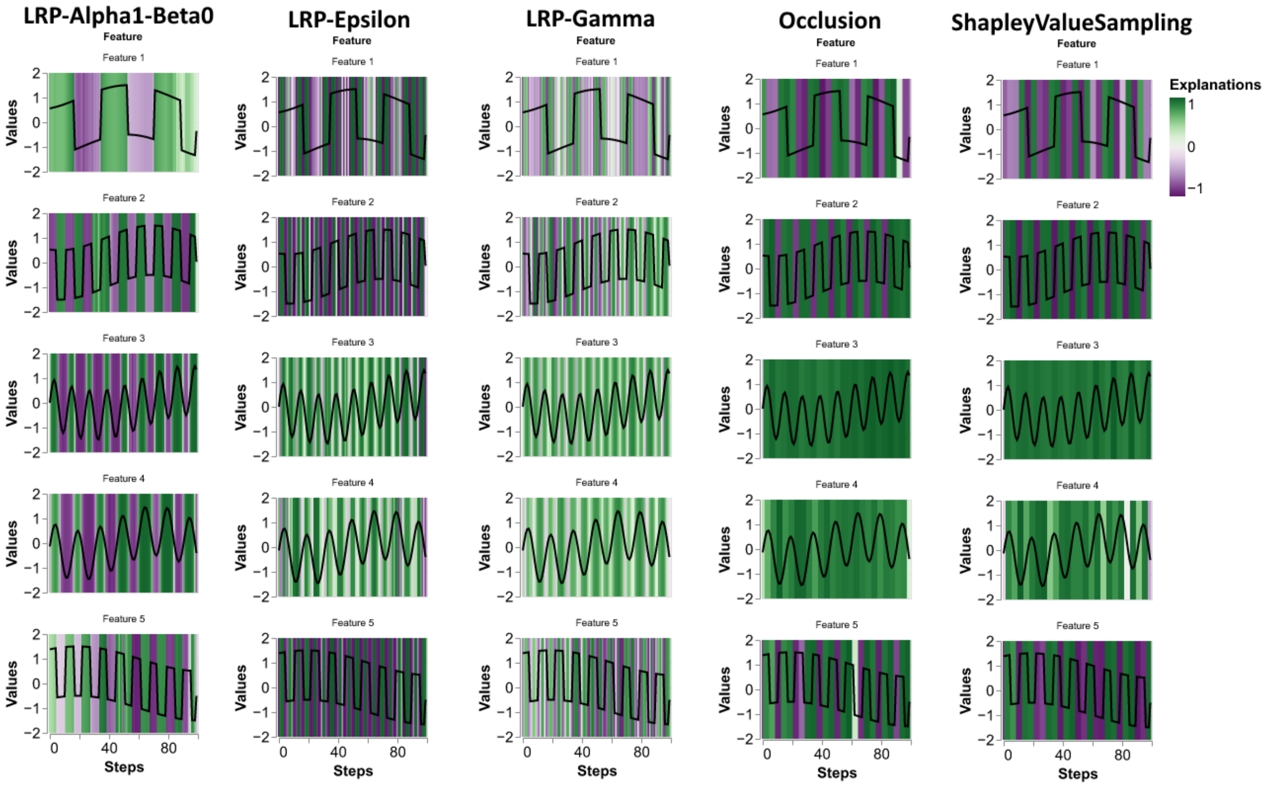


Figure S14: Median relevance attributions over all 100 iterations for one random sample for all XAI methods (Part 3).

| | MA | HE | EX | SE | OD | Normal |
|---|---|---|---|---|---|---|
| Deconvolution | -0.736 ±0.785 | -0.193 ±1.199 | 0.579 ±1.415 | 0.188 ±1.094 | 0.072 ±0.797 | 0.003 ±0.819 |
| InputXGradient | 0.395 ±1.356 | 0.007 ±0.945 | -0.312 ±1.191 | -0.005 ±0.966 | 0.011 ±0.708 | 0.0 ±0.7 |
| GradientShap | 0.358 ±1.319 | -0.002 ±0.896 | -0.329 ±1.347 | -0.012 ±0.947 | 0.006 ±0.797 | 0.0 ±0.693 |
| IntegratedGradients | 0.369 ±1.369 | -0.0 ±0.926 | -0.339 ±1.386 | -0.015 ±0.97 | 0.007 ±0.81 | 0.0 ±0.7 |
| Occlusion | -0.582 ±0.29 | -0.615 ±0.295 | -0.451 ±0.21 | -0.557 ±0.294 | -0.54 ±0.197 | -0.48 ±0.709 |
| KernelShap | -0.307 ±0.096 | -0.304 ±0.098 | -0.314 ±0.092 | -0.284 ±0.1 | -0.291 ±0.103 | -0.29 ±0.102 |
| DeepLiftShap | 0.429 ±1.406 | 0.005 ±0.955 | -0.365 ±1.318 | -0.303 ±0.986 | -0.029 ±0.756 | 0.0 ±0.713 |
| DeepLift | 0.429 ±1.406 | 0.005 ±0.955 | -0.365 ±1.318 | -0.303 ±0.986 | -0.029 ±0.756 | 0.0 ±0.713 |
| ShapleyValueSampling | -0.076 ±0.779 | -0.055 ±0.775 | -0.291 ±0.778 | -0.273 ±0.803 | -0.25 ±0.787 | 0.0 ±0.654 |
| LRP-Alpha1-Beta0 | 0.732 ±0.423 | 0.565 ±0.484 | 0.915 ±0.197 | 0.775 ±0.433 | 0.681 ±0.263 | 0.107 ±0.569 |
| LRP-Epsilon | 0.37 ±1.144 | 0.007 ±0.848 | -0.279 ±0.917 | -0.003 ±0.867 | 0.011 ±0.714 | 0.0 ±0.688 |
| LRP | 0.361 ±1.084 | 0.006 ±0.824 | -0.124 ±0.874 | -0.006 ±0.84 | 0.009 ±0.708 | 0.0 ±0.678 |
| GuidedBackprop | -0.751 ±1.421 | -0.272 ±1.331 | 0.709 ±1.722 | 0.469 ±1.325 | 0.369 ±1.09 | -0.025 ±0.886 |

Table S6: Median and interquartile ranges (IQR) for all correct classified class 1 samples over all 100 iterations for each abnormality. Abnormalities are Microaneurysms (MA), Haemorrhages (HE), Hard Exudates (EX), Soft Exudates (SE), Optic Disc (OD), and Normal for everything outside of the segmentation masks. LRP-$\gamma$ was removed from the table, because it did not converge for this dataset. Lime was removed from this table, because median and IQR were 0 for everything.
